# Identification of co-expressed gene sets for major molecular initiating events in rodent liver carcinogenesis using wild type and knock out rats

**DOI:** 10.64898/2026.09.15.751815

**Authors:** Kaitlyn K. Venneman, Manasi Kotulkar, Diego Paine-Cabrera, Jie Liu, Constance A. Mitchell, Keith Q. Tanis, J Christopher Corton, Udayan Apte

## Abstract

Liver tumors are the most common carcinogenic outcome observed in two-year bioassays used for regulatory risk assessment of chemicals and pharmaceuticals. Because these traditional long-term studies are resource-intensive and not feasible for most chemicals, there is a critical need to develop alternative approaches that can identify potential rodent liver carcinogens in short-term studies (e.g., ≤ 1 month). Biomarkers associated with molecular initiating events (MIEs) of early rodent liver tumorigenesis may enable the prediction of two-year outcomes from such studies. Here, we identified transcriptional changes associated with six key initiators of rat liver tumorigenesis: five transcription factors (TFs) associated with non-genotoxic modes of action (AhR, PXR, CAR, PPARα, and ERα), and tumor suppressor protein p53, which is activated by diverse cellular stressors including genotoxicity. Three known chemical activators for each TF were administered orally once daily for five days at doses associated with tumorigenic responses in previous rat cancer bioassays. RNA-seq identified differentially expressed genes (DEGs) that were consistently regulated by at least two activators in wild-type but not in corresponding TF knockout animals. Gene set enrichment analysis of ranked differential expression results demonstrated enrichment of pathways reflecting established TF biology. When comparing the consensus DEGs for each TF to a library of chemically induced, liver-specific gene expression profiles, we found that in most cases the chemicals with the most similar profiles to the consensus lists were known to activate the corresponding TF. Collectively, these findings identified candidate gene sets that may have utility for monitoring the induction of carcinogenesis-associated MIEs in short-term studies.

## Introduction

Since its inception in 1976, the two-year rodent bioassay has served as an important tool in chemical carcinogenicity testing (McConnell 1995). Agencies such as the U.S. Environmental Protection Agency (EPA) and Food and Drug Administration (FDA) have required data from these assays to support carcinogenicity risk assessments. However, the effectiveness of the two-year bioassay has long been questioned, and a growing movement now calls for reevaluation based on numerous scientific, ethical, and practical concerns (Cohen 2010; Doe et al. 2019; Goodman 2018; Stevens and Baker 2009; Suarez-Torres, Orozco, and Ciangherotti 2021). Within this context, the International Council for Harmonisation (ICH) released S1B(R1), which allows an integrated weight-of-evidence (WoE) approach to assess human carcinogenic risk for certain pharmaceuticals in lieu of a two-year rat study (ICH 2022). Additionally, the Rethinking Carcinogenicity Assessment for Agrochemical Projects (ReCAAP) framework provides a structured WoE approach for carcinogenicity assessment and, when justified, supports waivers for long-term rodent bioassays. This framework is increasingly being incorporated into regulatory risk assessment (EPA 2025; Hilton et al. 2022). However, fully replacing the two-year bioassay with more resource-efficient tools, such as *in vitro* systems, remains challenging (Jadhav, Gurgude, and Sawant 2025; Schmeisser et al. 2023). Short-term *in vivo* studies that measure the major early drivers of chemical-induced cancers can provide valuable insight into underlying mechanisms of toxicity (Cohen 2004; Hilton et al. 2022) and support the WoE used to assess whether a two-year study is of value. Such studies could then reduce reliance on two-year bioassays while serving as a bridge to shorter, mechanism-focused studies with greater translational relevance.

As the primary organ responsible for xenobiotic metabolism, the liver is particularly susceptible to chemical exposures. In addition, liver tumors are the most common carcinogenicity finding in two-year rodent bioassays used in risk assessment. This makes it an ideal candidate for studying carcinogenic risk (Gijbels and Vinken 2017; Jaeschke 2019; Rusyn et al. 2021; Walesky et al. 2013; Zhang, Qi, et al. 2022). As a result, numerous pathways that initiate liver tumorigenesis have been characterized. Events within these pathways can be organized by mode of action and adverse outcome to provide context for organ-specific toxicity and support development of novel tools and approaches to assess the potential and/or relevance of rodent bioassay findings. One such approach integrates toxicogenomic data into adverse outcome pathway-based frameworks to develop gene expression signatures, or biomarkers, offering a more quantitative approach for assessing carcinogenicity (Corton et al. 2022; Glaab et al. 2021; Podtelezhnikov et al. 2020; Rooney, Hill, et al. 2018).

The Health and Environmental Sciences Institute (HESI Global) Carcinogenomics Workgroup seeks to develop and validate genomic biomarkers that can be leveraged in short-term rodent studies to provide mechanism-based WoE and assess the utility of conducting a two-year bioassay. To do so, the Workgroup focuses on known MIEs that drive or sense adverse outcomes in rat liver (Corton et al. 2022). Included are the activation of xenobiotic receptors Aryl Hydrocarbon Receptor (AhR), Pregnane X Receptor (PXR), Constitutive Androstane Receptor (CAR), Peroxisome Proliferator-Activated Receptor Alpha (PPARα), and Estrogen Receptor Alpha (ERα), which are known to be non-genotoxic drivers of liver tumors in rats. Also included is the tumor suppressor protein p53, which is activated by diverse cellular stressors, including metabolic and oxidative stress, DNA damage, and hypoxia (Hernández Borrero and El-Deiry 2021). As such, biomarkers of p53 activation are often used as sentinels for potential genotoxicity, although follow-up investigation is required to determine the specific MIE and relevance to carcinogenicity.

As part of this larger initiative to develop and validate carcinogenesis-related biomarkers, short-term *in vivo* studies were conducted to identify transcriptomic changes after the activation of each MIE-associated TF (**Figure 1a-b**). By comparing transcriptional responses in wild-type rats to rats in which both alleles of the respective TF gene have been knocked out (TF-null rats), our strategy allowed us to unequivocally identify genes whose expression changes are dependent on the MIE-associated TF. The findings support the development and adjudication of TF-specific activity biomarkers in the rat liver.

**Figure 1:**
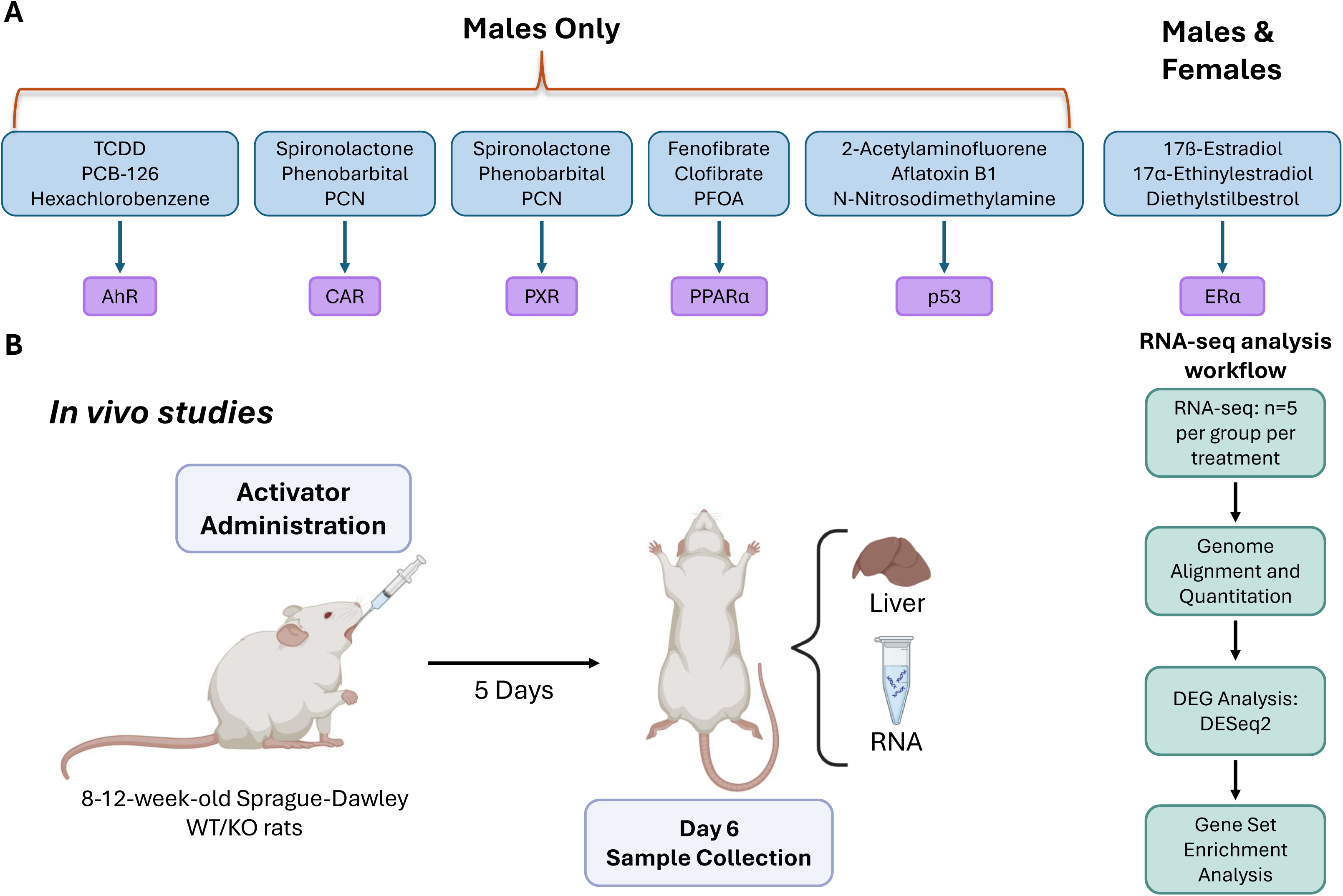
(A) Experimental protocol and (B) overall study design.

## Materials and Methods

### Animals

All animals were housed in facilities accredited by the Association for Assessment and Accreditation of Laboratory Animal Care at the University of Kansas Medical Center (KUMC) under a standard 12 hr light/dark cycle with free access to chow and water. All studies were approved by the Institutional Animal Care and Use Committee at KUMC. Sprague-Dawley male wild type (WT), AhR (HsdSage:SD-*Ahr^em1Sage^*), PXR (HsdSage:SD-*Nr1i2^tm1Sage^*), CAR (HsdSage:SD-*Nr1i3^tm1Sage^*), PPARα (HsdSage:SD-*Ppara^em1Sage^*), and p53 (HsdSage:SD:*Tp53^tm1Sage^*) knockout (KO) rats were purchased from Inotiv, Inc. (Lafayette, IN). These rats were originally developed at SAGE Labs, Inc., in St. Louis, MO and contain the truncated form of either the ligand binding domain (PPARα) or the DNA binding domain (other TFs), rendering them nonfunctional. The generation, genotyping, and characterization of the estrogen receptor alpha (*Esr1)*-knockout rats have been described previously in detail (Rumi et al. 2014). The breeding pairs used to generate rats for the *Esr1* arm of the study were provided by the lab of Dr. Soares at the University of Kansas Medical Center, and both male and female rats were used. An n = 5 was used for all groups, and all rats were aged 12-16 weeks at the time of treatment.

### Compound Exposure and Sample Collection

Compounds and doses for each group can be found in **Table 1**, and the experimental protocol is depicted in **Figure 1**. For the CAR and PXR studies, the activators spironolactone, phenobarbital, and PCN were chosen to investigate demonstrated crossover in transcriptomes as PCN preferentially activates PXR, phenobarbital preferentially activates CAR, and literature in mice indicated that spironolactone activates both TFs. All control WT and KO rats received an equivalent volume of vehicle, carboxymethylcellulose sodium (CMC NA, Sigma-Aldrich cat# C5678). Chemical selection and dosing were based on previous rat *in vivo* studies (**Table 1**) and were focused on identifying a dose expected to induce tumorigenesis in a two-year bioassay or, if tumorigenic doses were not known, shown to induce a sufficient level of gene expression in a shorter-term rat *in vivo* study (e.g., PPARα: *Cyp4a10* for PFOA or CAR: *Cyp2b10* for phenobarbital). Rats were treated via oral gavage once daily for five days and euthanized on the sixth day approximately 24 hrs after the final dose was given. At the time of necropsy, blood was collected and allowed to clot at room temperature for 10min and then centrifuged at 7000 rcf for 15 min at 4° C to isolate serum. Livers were removed and weighed to calculate liver to bodyweight (LW:BW) ratios. A portion of the liver was fixed in 4% formaldehyde for 48 hrs, followed by an additional 24 hrs in ethanol. These were then processed to obtain paraffin-embedded tissue sections for histology. A second portion of the liver was cryopreserved in O.C.T compound (Fisher Scientific cat# 23-730-571). All remaining liver tissue was flash-frozen in liquid nitrogen and stored at -80 °C for future analysis. In addition to the liver, several other organs including bone, fat, ileum, muscle, kidney, pancreas, testes, and ovaries (ERα females only) were also collected, flash-frozen in liquid nitrogen, and stored at –80 °C. Only liver tissues were used in the present study.

**Table 1:** Study groups and their corresponding compounds. Doses for each compound are listed as well as the vendor from where the compound was acquired.

| Group | Compound | Dose | Reference | Vendor |
| --- | --- | --- | --- | --- |
| AhR | 2,3,7,8-tetrachlorodioxin (TCDD) | 150 ng/kg | (Qin et al. 2019)**<br>(Goodman and Sauer 1992)* | AccuStandard cat# D-404S |
|  | PCB-126 | 1000 ng/kg | National Toxicology Program;<br>(Qin et al. 2019)* | Iowa Superfund Research Program (ISRP) Synthesis Core |
|  | Hexachlorobenzene | 100 mg/kg | IARC (2001). Monograph Vol. 79, Hexachlorobenzene*<br>(Qin et al. 2019)** | Sigma Aldrich cat# 45522 |
| CAR/PXR | Spironolactone | 300 mg/kg | IARC (2001). Monograph Vol. 79, Spironolactone* | Sigma Aldrich cat# S3378+ |
|  | Phenobarbital | 500 mg/kg | IARC (2001). Monograph Vol. 79, Phenobarbital* | KUMC LAR |
|  | Pregnenolone Carbonitrile (PCN) | 125 mg/kg | (Guzelian et al. 2006)**<br>(Forbes et al. 2017)** | Cayman Chemical cat# 16343 |
| PPARα | Fenofibrate | 200 mg/kg | FDA Label<br>Drugbank: Fenofibric Acid | Sigma-Aldrich cat# F6020 |
|  | Clofibrate | 500 mg/kg | IARC (1980). Monograph Vol. 24 | TCI cat# C0941 |
|  | PFOA | 5 mg/kg | (Cui et al. 2009)** | Sigma-Aldrich cat# 77262 |
| p53 | 2-Acetylaminofluorene (2-AAF) | 30,000 μg/kg | (Weisburger et al. 1981)* | Sigma Aldrich cat# 8205760010 |
| | Aflatoxin B1 (AFB1) | 300 μg/kg | (Wogan, Paglialunga, and Newberne 1974)* | Sigma-Aldrich cat# A6636 |
| | N-Nitrosodimethylamine (NDMA) | 300 μg/kg | (Peto et al. 1991)* | Supelco Analytics cat# PHR2407 |
| ERα | 17β-Estradiol | 10 mg/kg | National Toxicology Program;<br>(Williams et al. 1993) | Sigma Aldrich cat# 3301 |
| | 17α-Ethinylestradiol | 10<br>mg/kg | National Toxicology Program | Sigma<br>Aldrich cat#<br>E4876 |
|  | Diethylstilbestrol<br>(DES) | 10<br>mg/kg | (Williams et al. 1993)* | Sigma<br>Aldrich cat#<br>D4628 |

\*Selected based on cancer bioassay in rats
\*\*Selected based on gene expression study in rats

### RNA-Sequencing

Total RNA was isolated from frozen liver tissue portions using the MagMAX™-96 for Microarrays Total RNA Isolation Kit (ThermoFisher AM1839). For each sample, the liver was harvested from processed samples using the MagMAX™ Express-96 Deep Well Magnetic Particle Processor (ThermoFisher) as per the manufacturer’s instructions. Total RNA was quantified via NanoDrop (ThermoFisher ND8000) and 500ng was used per sample for library preparation using the KAPA mRNA HyperPrep Kit (Roche 08098123702) as per the manufacturer’s instructions. Sequencing was performed on the Illumina NextSeq2000 using the NextSeq 2000 P3 Reagents (50 Cycles) (Illumina 20046810) or the NovaSeqX Platform using NovaSeq X Series 10B Reagent Kit (100 cycles) (Illumina 20085596). Genome alignment and gene quantitation were performed using OmicSoft Array Studio. Reads were aligned to the Rat.B7.2 genome reference using the OmicSoft Aligner with a maximum of 2 allowed mismatches. Gene level counts were determined by the RNA-Seq expectation maximization algorithm as implemented in OmicSoft Array Studio and using RefSeq_RS_2023_06 gene models. Ratio values for each gene were calculated as the log_10_ (FPKM (fragments per kilobase per million fragments) + 0.001) for each animal minus the average log_10_ (FPKM + 0.001) of the concurrent animals.

### Bioinformatic Analysis

Total original gene counts of RNA sequencing were imported into the Partek Flow Server. Differentially expressed gene (DEG) analysis was performed with the DESeq2 method using the pair-fed group as the control. WT treated rats were then compared to their corresponding KO treated rats to identify DEGs unique to WT treatment (p-value < 0.05, |log_2_FC| > 1). Because these are short-term studies and some treatments expectedly showed weaker responses, this comparison was repeated at a less stringent threshold (p-value 0.05 only) to capture more subtle changes. Gene Set Enrichment Analysis (GSEA) was performed using the fgsea R package with the Hallmark gene sets for *Rattus norvegicus* from MSigDB (Liberzon et al. 2015) to identify common pathways between treatments. In GSEA, genes were ranked according to the Wald statistic from DESeq2, which measures the strength of differential expression. Enrichment scores were calculated with 10,000 permutations (nperm = 10000) to estimate significance, and ties in the ranking were handled arbitrarily by fgsea. Pathways with FDR < 0.05 were considered significant. Uncharacterized loci (LOC-designated transcripts) were identified and excluded from GSEA analyses due to limited functional annotation. Complete differential expression results, including LOCs, are provided in **Files S1a-S1g**.

### Consensus Gene Lists

First, receptor-specific gene expression signatures were built by contrasting gene expression changes in the WT rats to changes in the KO rats. Changes specific to WT rats were then compared to a library of chemically induced, liver-specific gene expression profiles to determine the final consensus lists. For the analysis of PXR, PPARα, p53, ERα males, and ERα females, the transcript profiles from all three chemicals in WT and KO rats were used. Given that hexachlorobenzene (HCB) was a weak AhR activator, only TCDD and PCB-126 profiles were used to generate the AhR consensus gene list. As phenobarbital was the only preferential CAR activator that was used in the analysis, its consensus gene list was generated from a comparison of phenobarbital treatment in WT vs. CAR KO rats. Starting with the statistically filtered gene lists (uncorrected p-value < 0.05), genes were selected if they exhibited consistent directional changes between the chemicals in the WT rats and no changes or opposite changes in the KO rats. Since the size of gene lists can have an impact on the significance of the overlap in pair-wise comparisons, gene lists over 100 genes were filtered to include only 100 or fewer genes for consistency. This was carried out on the lists derived from ERα males, ERα females, and CAR studies to remove any gene with an absolute fold-change less than 2.5-fold. The final number of genes after import into BaseSpace Correlation Engine (Illumina; BSCE) were 17 (AhR), 68 (PXR), 51 (CAR), 77 (PPARα), 43 (p53), 91 (ERα males), and 71 (ERα females). There were 19 genes that overlapped between the ERα male and female lists. Minimal overlap occurred between other gene sets (**File S2a**). A total of 370 genes were identified in the consensus gene sets.

### Graphs and Statistical Analysis

Heat maps were produced in RStudio (Version 4.3.3, RStudio Team) using the R packages ggplot2 (Version 3.5.1) and pheatmap (Version 1.0.13). For PXR, CAR, PPARα, p53, and ERα heat maps, the top 15 upregulated and top 15 downregulated DEGs were selected based on log_2_FC. This was not the case for AhR as there were only 29 DEGs identified. Venn diagrams were generated using the online tool Venny 2.1.0 (http://bioinfogp.cnb.csic.es/tools/venny/). Bar graphs and their corresponding statistical analyses were produced in GraphPad Prism 8. Since more than three groups were compared in the statistical analysis for LW:BW ratio, an ANOVA was used followed by a Tukey multiple comparison post-hoc test. Statistical significance was considered when the p-value was < 0.05.

## Results

### Transcriptomic effects of Aryl Hydrocarbon Receptor activation

WT and AHR KO rats were treated with either TCDD, PCB-126, or hexachlorobenzene (HCB). We identified 27 DEGs in WT TCDD-treated rats, none of which were identified in KO treated rats. In WT PCB-treated rats, we identified 36 DEGs, of which 33 (92%) were not identified in the KO. In WT HCB-treated rats, we identified 16 DEGs, of which 13 (81%) were not identified in the KO (p-value < 0.05, |log_2_FC| > 1) (**Supplemental Figures 2a-c**). Additional genes were also identified using a significance cutoff of p-value < 0.05 only (**Supplemental Figures 2d-f**). We then compared the WT DEGs for each treatment (p-value < 0.05) and identified 29 common genes between two or more of the three treatments (**Figure 2a**). Comparison of WT and KO expression profiles revealed that these treatment-responsive genes in WT showed a weaker or absent response in KO (**Figure 2b**). Common significant pathways (FDR < 0.05) identified with GSEA are shown in **Figure 2c** and include pathways consistent with known biology, such as xenobiotic metabolism. Liver to body weight (LW:BW) ratio was unaffected by treatment with TCDD, PCB-126, or HCB (**Supplemental Figure 1a**).

**Figure 2:**
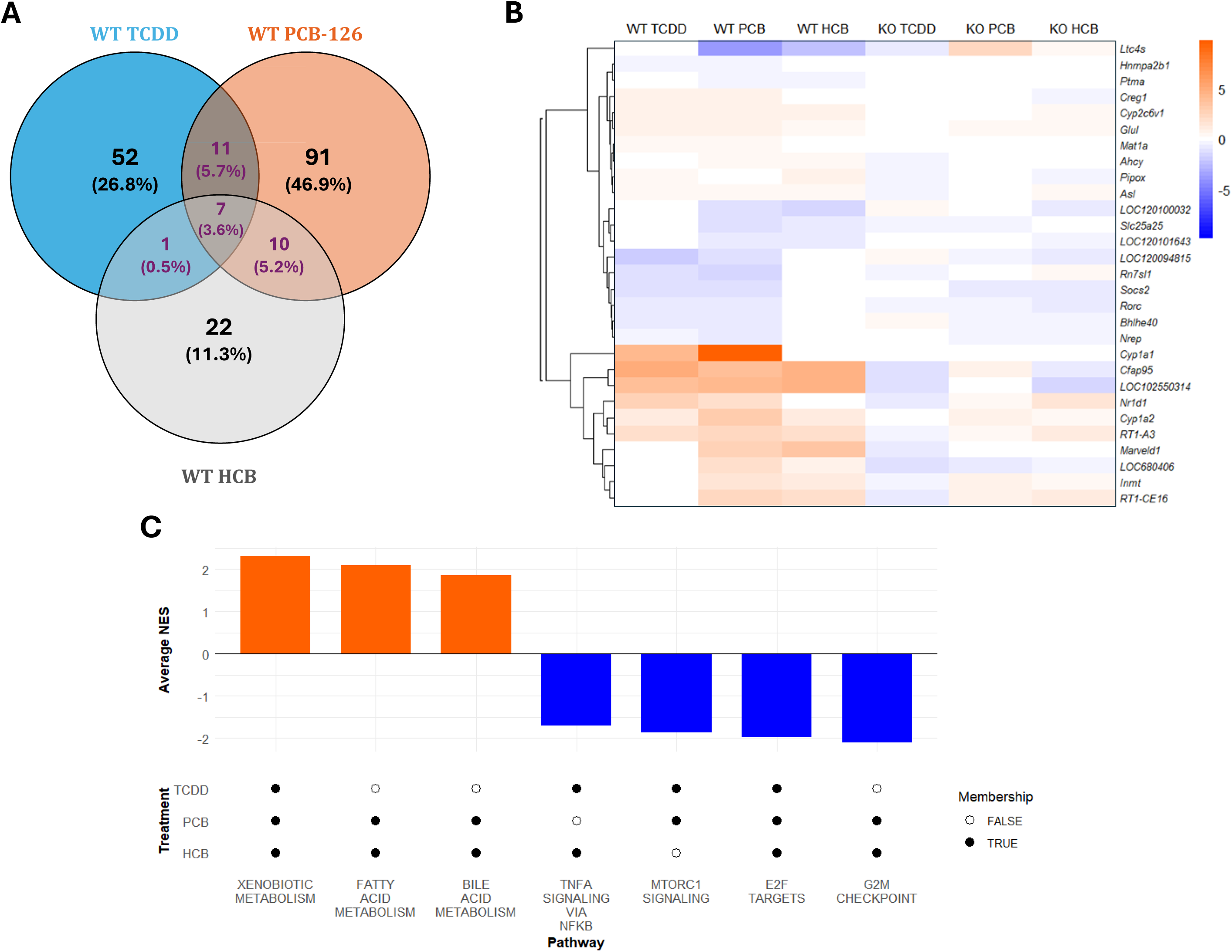
Transcriptomic changes after activator exposure in AhR study. (A) Venn diagram showing number of DEGs in WT after treatment with either TCDD, PCB-126, or HCB. Highlighted numbers indicate genes common between 2 or more of the 3 treatments (p-value < 0.05). (B) Heatmap of common DEGs highlighting shared genes in 2 or more of the 3 treatments. (C) Bar graph showing common pathways in 2 or more of the 3 treatments (FDR < 0.05). Average NES was calculated using NES from compounds that have membership.

### Transcriptomic effects of Pregnane X Receptor activation

We identified 91 DEGs in WT spironolactone treated rats, 85 (93%) of which were not identified in PXR KO rats. Of the 38 DEGs identified in WT PCN treated rats, 37 (97%) were not identified in the KO rats. In contrast, 38 (52%) of the 73 DEGs identified in WT phenobarbital treated rats were also identified in the PXR KO rats (p-value < 0.05 and |log_2_FC| > 1) (**Supplemental Figures 3a-c**). This is consistent with phenobarbital being predominantly a CAR agonist in rats (Podtelezhnikov 2020). Additional genes identified using p-value < 0.05 only are shown in **Supplemental Figures 3d-f**. Comparing WT DEGs for each treatment identified 79 genes common in two or more of the three treatments (**Figure 3a**). The top 30 up and downregulated DEGs, selected as described in the Methods, are depicted in **Figure 3b**. Comparison of WT and KO expression profiles revealed that most treatment-responsive genes in WT showed a weaker or absent response in KO. Top common significant pathways (FDR < 0.05) identified with GSEA are shown in **Figure 3c** and include pathways consistent with known biology, such as xenobiotic and fatty acid metabolism. In addition, there were significant increases in the LW:BW ratio in WT rats treated with spironolactone and PCN. This effect was not seen in WT rats treated with phenobarbital or KO rats after any treatment (**Supplemental Figure 1b**).

**Figure 3:**
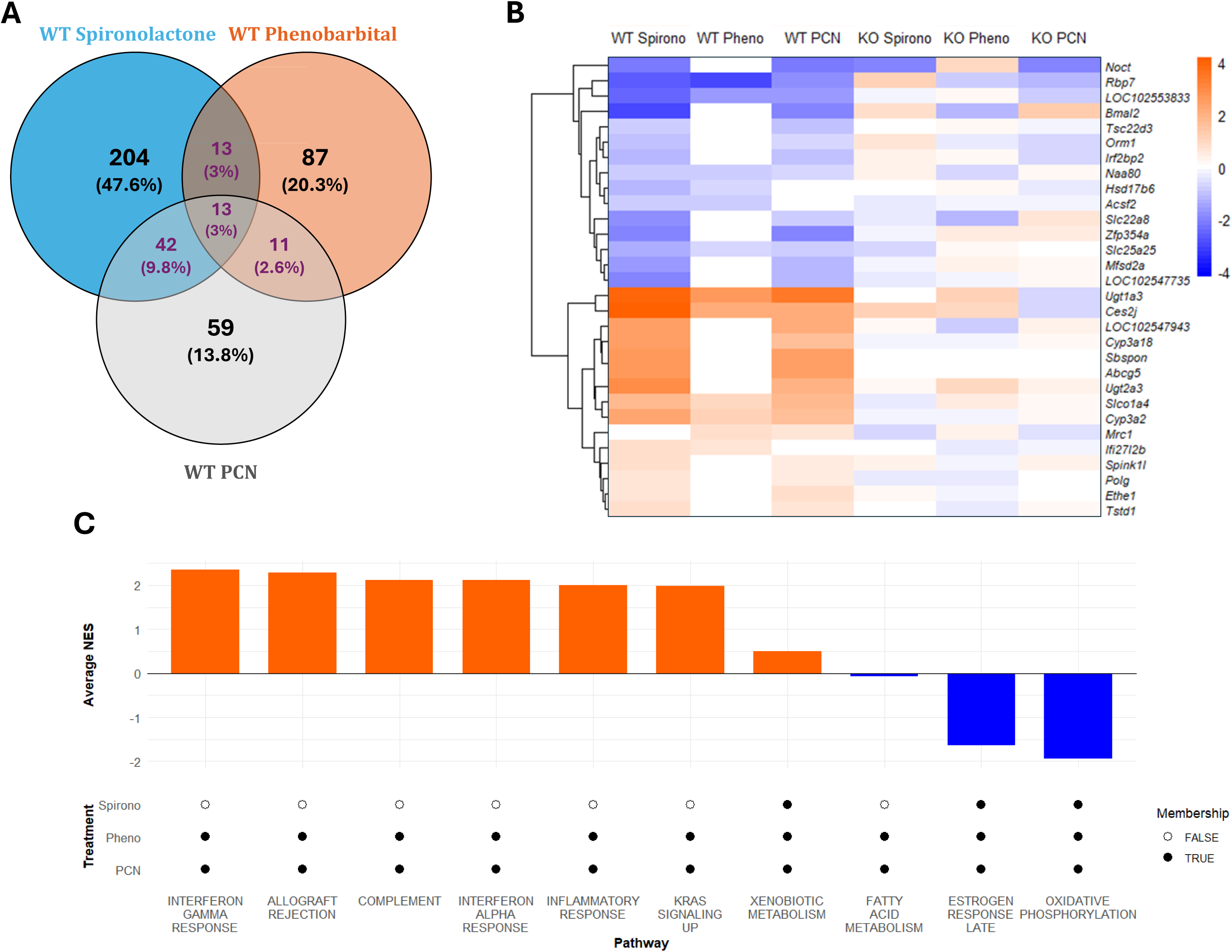
Transcriptomic changes after activator exposure in PXR study. (A) Venn diagram showing number of DEGs in WT after treatment with either Spironolactone, Phenobarbital, or PCN. Highlighted numbers indicate genes common between 2 or more of the 3 treatments (p-value < 0.05). (B) Heatmap of top 30 common DEGs highlighting shared genes in 2 or more of the 3 treatments. (C) Heatmap comparing WT to KO after treatment with phenobarbital only. (D) Bar graph showing top 10 common pathways in 2 or more of the 3 treatments (FDR < 0.05). Average NES was calculated using NES from compounds that have membership.

### Transcriptomic effects of Constitutive Androstane Receptor activation

In a separate cohort, WT and CAR KO rats were also treated with either spironolactone, phenobarbital, or PCN. Comparison identified 83 DEGs for WT phenobarbital treated rats, of which 78 (94%) were not identified in CAR KOs. In contrast, of the 208 DEGs identified for WT spironolactone treated rats, 87 (42%) were still observed in CAR KOs. Of the 51 DEGs identified in WT PCN treated rats, 17 (33%) were still observed in the CAR KOs (p-value < 0.05, |log_2_FC| > 1) (**Supplemental Figures 4a-c**). This is consistent with the known PXR activity of PCN and spironolactone. In this case, comparing the WT DEGs for each treatment revealed 84 genes common between two or more of the three treatments (**Figure 4a**). The top 30 up and downregulated DEGs are depicted in **Figure 4b**. Here too, comparison of WT and KO expression profiles revealed that treatment-responsive genes in WT showed a weaker or absent response in KO. Since both PCN and spironolactone proved to be highly preferential for PXR, **Figure 4c** depicts the top 30 up and down regulated DEGs after phenobarbital treatment only. Consistent with **Figure 4b**, comparison of WT and KO expression profiles revealed that treatment-responsive genes in WT showed a weaker or absent response in KO. Additional genes identified using p-value < 0.05 only are shown in **Supplemental Figures 4d-f**. Top common significant pathways (FDR < 0.05) identified with GSEA are shown in **Figure 4d** and include pathways consistent with known biology, such as xenobiotic and fatty acid metabolism. As with the AhR studies, LW:BW ratio was unaffected by treatment with spironolactone, phenobarbital, or PCN (**Supplemental Figure 1c**).

**Figure 4:**
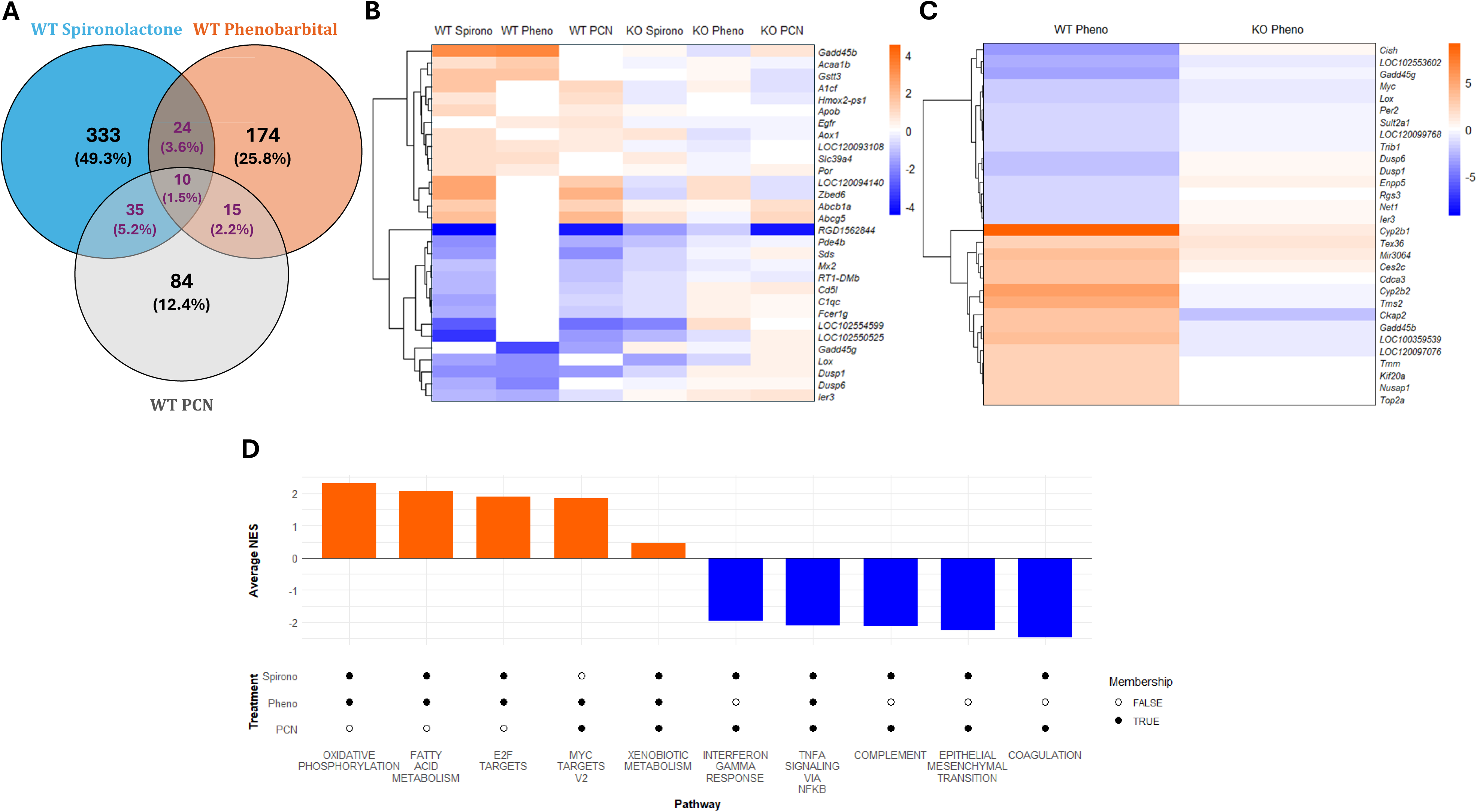
Transcriptomic changes after activator exposure in CAR study. (A) Venn diagram showing number of DEGs in WT after treatment with either Spironolactone, Phenobarbital, or PCN. Highlighted numbers indicate genes common between 2 or more of the 3 treatments (p-value < 0.05). (B) Heatmap of top 30 common DEGs highlighting shared genes in 2 or more of the 3 treatments. (C) Heatmap of top 30 DEGs in WT after phenobarbital treatment. (D) Bar graph showing top 10 common pathways in 2 or more of the 3 treatments (FDR < 0.05). Average NES was calculated using NES from compounds that have membership.

### Transcriptomic effects of Peroxisome Proliferator-Activated Receptor Alpha activation

WT and PPARα KO rats were treated with fenofibrate, clofibrate, or PFOA. In WT rats treated with fenofibrate, we identified 29 DEGs, of which 28 (97%) were not identified in the KO rats. We identified 492 DEGs in WT clofibrate treated rats, of which 475 (97%) were not identified in the KO rats. Of the 458 DEGs identified in WT PFOA treated rats, 426 (93%) were not identified in the KO rats (p-value < 0.05, |log_2_FC| > 1) (**Supplemental Figures 5a-c**). Additional genes identified using p-value < 0.05 only are shown in **Supplemental Figures 5d-f**. Comparison of WT DEGs for these treatments revealed 1,096 common genes in two or more of the three treatments (**Figure 5a**). The top 30 up and downregulated DEGs are shown in **Figure 5b**. As with the other receptors, comparison of WT and KO expression profiles revealed that most treatment-responsive genes in WT showed a weaker or absent response in KO. Top common significant pathways (FDR < 0.05) identified with GSEA are shown in **Figure 5c** and include pathways consistent with known biology, such as fatty acid metabolism and peroxisome-related pathways. A significant increase in LW:BW ratio was only apparent in WT clofibrate treated rats (**Supplemental Figure 1d**).

**Figure 5:**
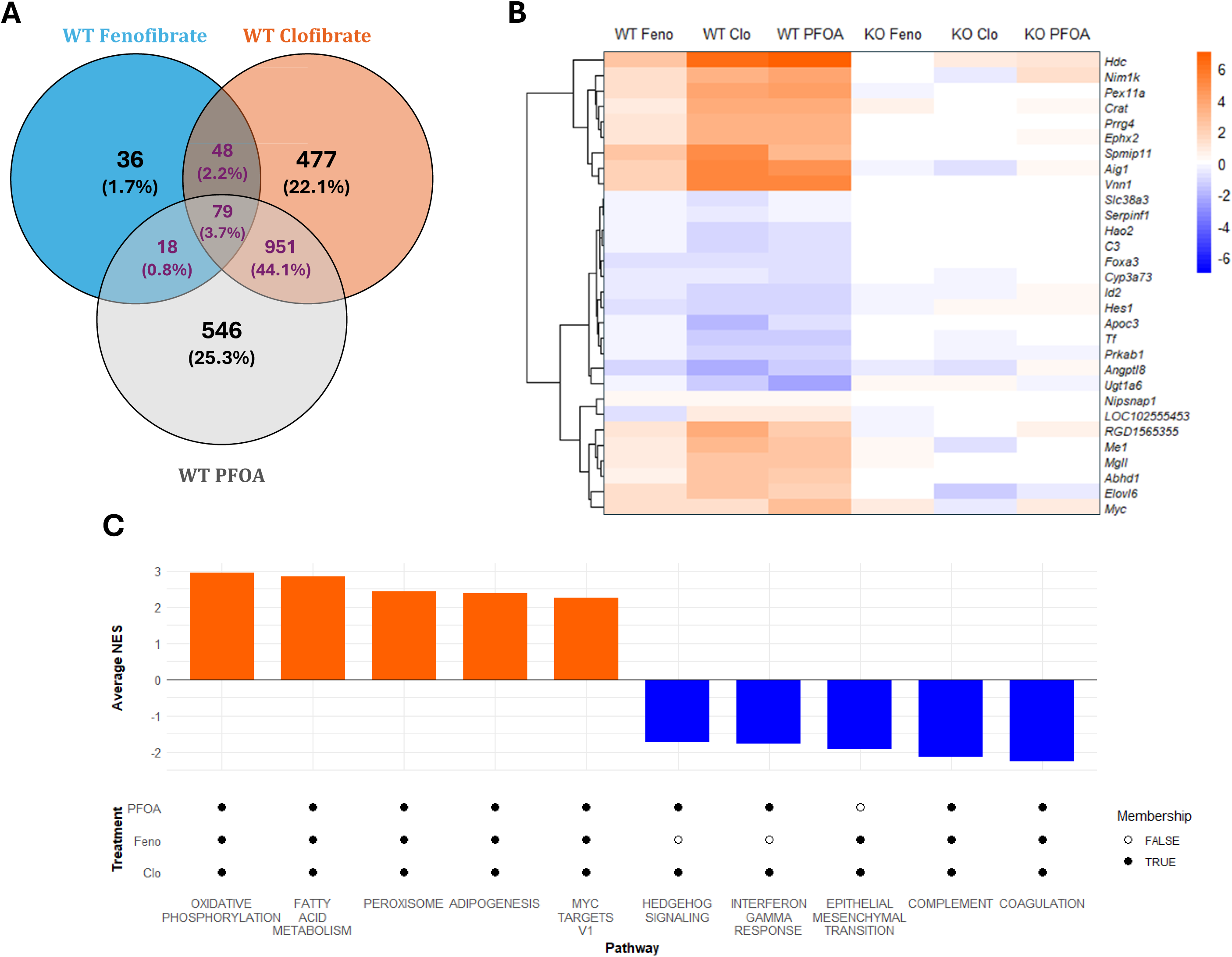
Transcriptomic changes after activator exposure in PPARα study. (A) Venn diagram showing number of DEGs in WT after treatment with either Fenofibrate, Clofibrate, or PFOA. Highlighted numbers indicate genes common between 2 or more of the 3 treatments (p-value < 0.05). (B) Heatmap of top 30 common DEGs highlighting shared genes in 2 or more of the 3 treatments. (C) Bar graph showing top 10 common pathways in 2 or more of the 3 treatments (FDR < 0.05). Average NES was calculated using NES from compounds that have membership.

### Transcriptomic effects of p53 activation

WT and p53 KO rats were treated with the genotoxicants 2-AAF, Aflatoxin B1 (AFB1), or n-nitrosodimethylamine (NDMA). Of the 264 DEGs identified in the WT 2-AAF treated rats, 245 (93%) were not identified in the KO treated rats. We identified 317 DEGs in the WT AFB1 treated rats, of which 294 (93%) were not identified in the KOs. For NDMA, we identified 50 DEGs in WT treated rats, 40 (80%) of which were not identified in KO treated rats. (p-value < 0.05, |log_2_FC| > 1) (**Supplemental Figures 6a-c**). Additional genes identified using p-value < 0.05 only are shown in **Supplemental Figures 6d-f**. Comparison of WT DEGs for these treatments revealed 515 common genes in two or more of the three treatments (**Figures 6a**). The top 30 up and downregulated DEGs are shown in **Figure 6b**. Once again, comparison of WT and KO expression profiles revealed that most treatment-responsive genes in WT showed a weaker or absent response in KO (**Figure 6b**). Top common significant pathways (FDR < 0.05) identified with GSEA are shown in **Figure 6c** and include pathways consistent p53 activation, such as the p53 pathway, apoptosis, and reactive oxygen species signaling. However, additional pathways were also represented, such as fatty acid metabolism. A significant increase in LW:BW ratio was only apparent in WT rats treated with NDMA (**Supplemental Figure 1e**).

**Figure 6:**
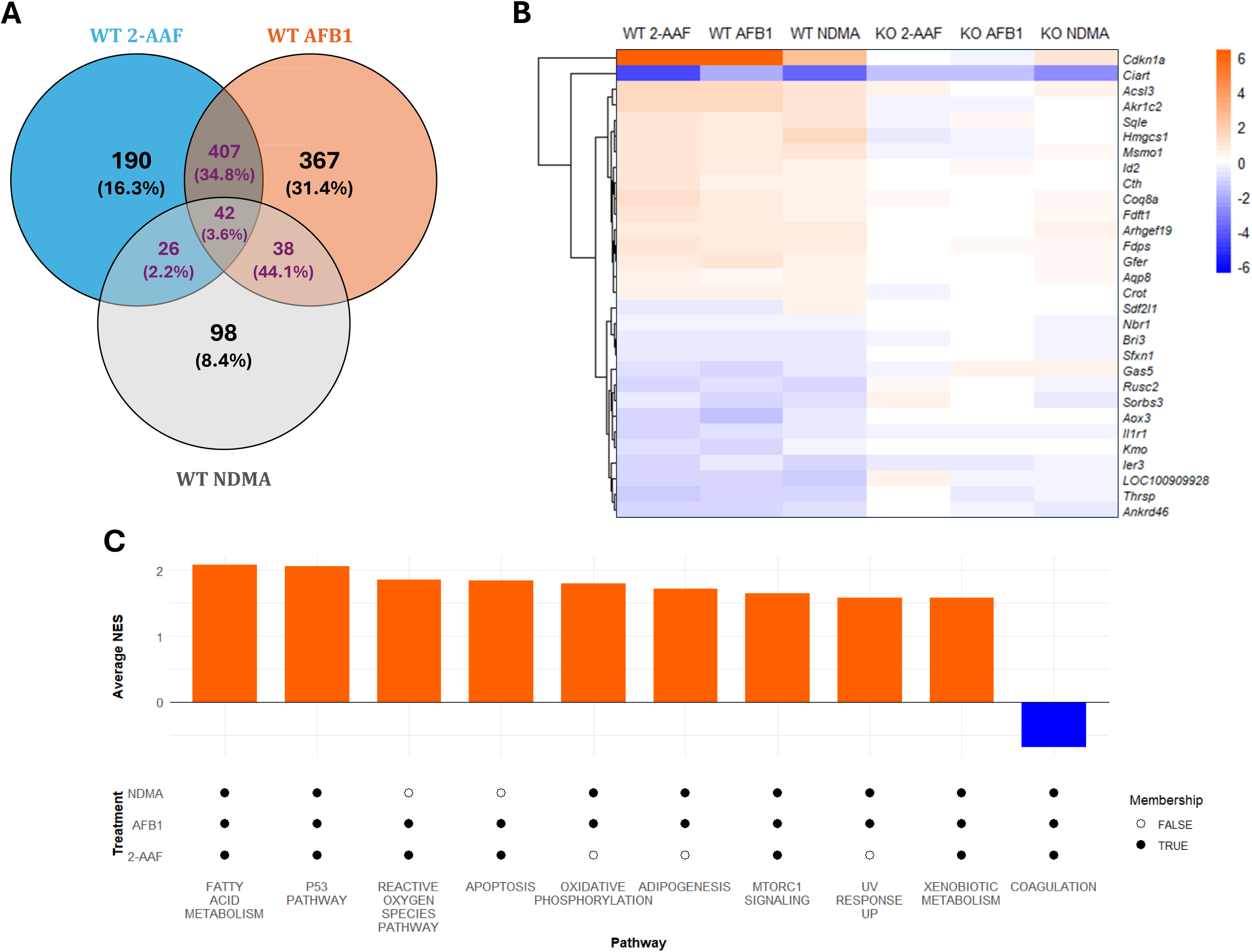
Transcriptomic changes after activator exposure in p53 study. (A) Venn diagram showing number of DEGs in WT after treatment with either 2-AAF, AFB1, or NDMA. Highlighted numbers indicate genes common between 2-3 out of 3 treatments (p-value < 0.05). (B) Heatmap of top 30 common DEGs highlighting shared genes in 2 or more of the 3 treatments. (C) Bar graph showing top 10 common pathways in 2 or more of the 3 treatments (FDR < 0.05). Average NES was calculated using NES from compounds that have membership.

### Transcriptomic effects of Estrogen Receptor Alpha activation

Male and female WT and ERα KO rats were treated with 17ß-estradiol (17ß), 17α-ethinylestradiol (17α), or diethylstilbestrol (DES). In WT males treated with 17ß, we identified 140 DEGs, of which 138 (99%) were not identified in the male KO treated rats. For WT 17α treated males, we identified 356 DEGs, 354 (99%) of which were not identified in male KOs. We identified 349 DEGs in WT DES treated males, of which 341 (98%) were not identified in male KOs. (p-value < 0.05, |log_2_FC| > 1) (**Supplemental Figures 7a-c**). In treated males, we identified 997 DEGs common between two or more of the three treatments (**Figure 7a**). The top 30 up and downregulated DEGs are shown in **Figure 7b**, and the reduced responsiveness of treatment-induced genes in the KO compared to the WT is even more apparent here compared to the other groups. Top common significant pathways (FDR < 0.05) identified with GSEA are shown in **Figure 7c** and include pathways consistent with known biology, such as fatty acid and bile acid metabolism. LW:BW ratio showed a significant increase in WT males treated with DES, an effect that was not seen in any other groups (**Supplemental Figure 1f**).

**Figure 7:**
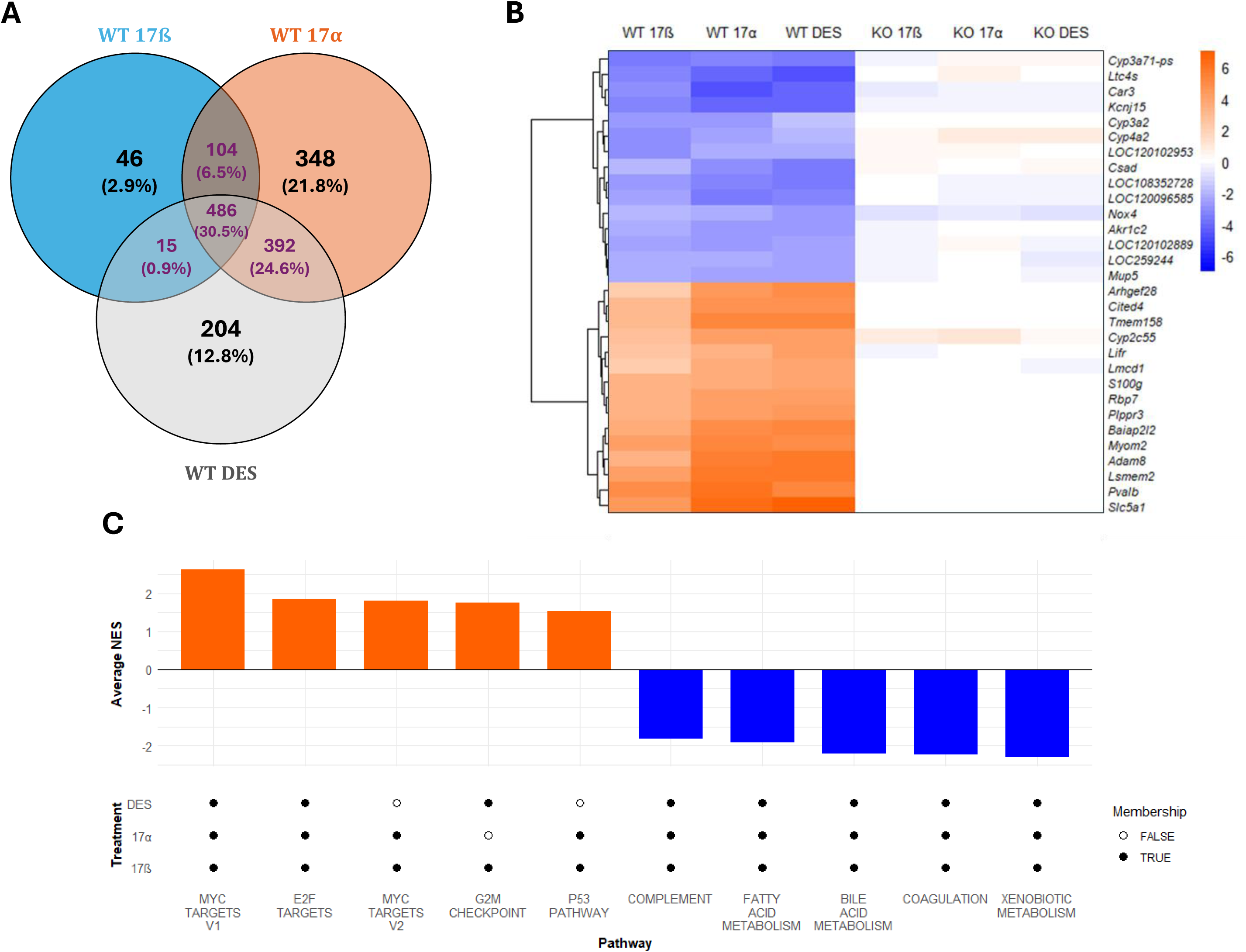
Transcriptomic changes after activator exposure in ERα male study. (A) Venn diagram showing number of DEGs in WT after treatment with either 17ß-estradiol, 17α-ethinylestradiol, or diethylstilbestrol. Highlighted numbers indicate genes common between 2 or more of the 3 treatments (p-value < 0.05). (B) Heatmap of top 30 common DEGs highlighting shared genes in 2 or more of the treatments. (C) Bar graph showing top 10 common pathways in 2 or more of the 3 treatments (FDR < 0.05). Average NES was calculated using NES from compounds that have membership.

In females treated with 17ß, we identified 160 DEGs, of which 158 (99%) were not identified in female treated KOs. For 17α, we identified 279 genes in female treated WT rats, none of which were identified in the KOs. Of the 237 genes identified in WT DES treated females, 236, only one DEG was shared with the KO (p-value < 0.05, |log_2_FC| > 1) (**Supplemental Figure 8a-c**). In treated females, we identified 866 genes common between two or more of the three treatments (**Figures 8a**). The top 30 up and downregulated DEGs are shown in **Figure 8b**. As with ERα males, the reduced responsiveness of treatment-induced genes in the KO compared to the WT is even more apparent here compared to the other groups (**Figure 8b**). Top common significant pathways (FDR < 0.05) identified with GSEA are shown in **Figure 8c** and also include pathways consistent with known biology, such as fatty acid and bile acid metabolism. In this case, LW:BW ratio showed an increase in WT females treated with 17α or DES, which was not seen on any other groups (**Supplemental Figure 1g**). Additional genes for WT and KO males and females for each treatment using a cutoff of p-value < 0.05 only are shown in **Supplemental Figures 7d-f** and **Supplemental Figures 8d-f**.

**Figure 8:**
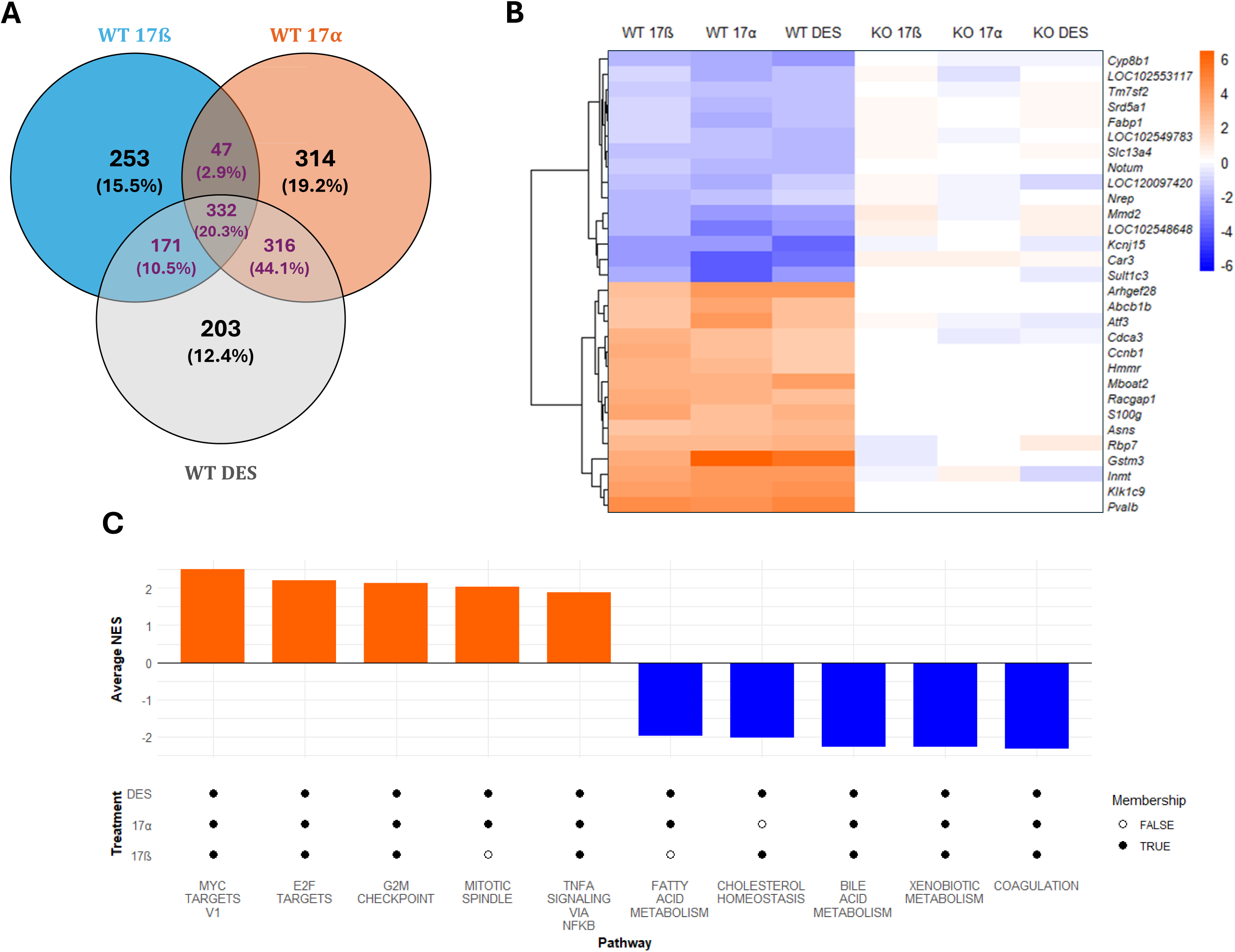
Transcriptomic changes after activator exposure in ERα female study. (A) Venn diagram showing number of DEGs in WT after treatment with either 17ß-estradiol, 17α-ethinylestradiol, or diethylstilbestrol. Highlighted numbers indicate genes common between 2 or more of the 3 treatments (p-value < 0.05). (B) Heatmap of top 30 common DEGs highlighting shared genes in 2 or more of the 3 treatments. (C) Bar graph showing top 10 common pathways in 2 or more of the 3 treatments (FDR < 0.05). Average NES was calculated using NES from compounds that have membership.

### Identification of TF activation using consensus gene sets

Transcript profiles were used to generate consensus gene sets for each TF as described in the Methods. Each gene set was compared to a compendium of transcript profiles derived from the livers of compound-treated rats to determine the ability of the gene set to identify compounds that activate the same TF. The -log(p-value) of the top 10 correlations between the consensus gene set and the compound vs. control comparisons for each MIE are shown in **Figure 9a-g**. **Figure 9h** provides a summary showing the number of known and unknown activators for each MIE. In the following analysis, we describe the top 10 profiles that exhibit the greatest correlations to the consensus gene lists (**Figure 9a-h; File S2b**).

- For AhR, existing literature demonstrates that leflunomide (Patel et al. 2015), oxfendazole (Dewa et al. 2007), and pantoprazole (Masubuchi and Okazaki 1997) activate AhR. Podtelezhnikov et al. identified gene sets that were used to assess the activation of 10 TFs in the livers of treated rats, including those examined in this study (Podtelezhnikov et al. 2020). There was an overlap between our predictions and the predictions for AhR in their study, which included erlotinib (both dose levels), miconazole, and rolipram, but not rosiglitazone, tamoxifen, and thioguanine (≥0.4 loading value from (Podtelezhnikov et al. 2020)). None of the top 10 hits included one of the many TCDD profiles in the compendium, and the significance of the correlations to the AhR consensus was relatively low compared to the other consensus gene sets described below. Together, this suggests additional refinement of the consensus set may be required for improved sensitivity and specificity.
- All of the compounds identified using the PXR consensus set are known PXR activators, including clotrimazole, “compound Z”, cyproterone acetate, econazole, fipronil, miconazole, mifepristone, and pregnenolone carbonitrile (AbdulHameed, Ippolito, and Wallqvist 2016; Ma, Idle, and Gonzalez 2008; Pinne, Ponce, and Raucy 2016; Podtelezhnikov et al. 2020; Roques et al. 2013).
- For CAR, all compounds are known or predicted CAR activators, including phenobarbital (Men and Wang 2023), artemisinin (Simonsson et al. 2006), clotrimazole (Marx-Stoelting, Knebel, and Braeuning 2020), alpidem, anastrozole, econazole, ethionamide, panadiplon, and promethazine (Podtelezhnikov et al. 2020).
- All of the compounds identified using the PPARα consensus set are well-known activators, including bezafibrate, clofibrate, dehydroepiandrosterone, fenofibrate, and WY-14,643 (Corton, Peters, and Klaunig 2018; Mastrocola et al. 2003).
- The p53 consensus gene set identified only three compounds with known ability to activate p53, including thioctic acid (Simbula et al. 2007), acetamidofluorene (Ohlson, Koroxenidou, and Hällström 1998), and ethionine (Tsujiuchi et al. 1997). Notably, of these three compounds, only acetamidofluorene is a direct genotoxicant, highlighting the diversity of p53 inputs. The other identified compounds, however, have not been previously linked to p53 activation and include gemfibrozil, lovastatin, and a chemical called “R1”. Gemfibrozil and lovastatin are used to lower cholesterol, and research shows both can activate SREBP2, leading to increases in cholesterol synthesis genes (Roglans et al. 2001; Sheng et al. 1995). In addition, the original study for “R1” describing liver gene expression did note increases in cholesterol synthesis genes (Roth et al. 2011). Furthermore, the p53 consensus gene set also included genes found in a mouse liver biomarker predictive of modulation of SREBP family members (TFs involved in cholesterol regulation) (Corton 2019). The overlapping genes were those involved in cholesterol biosynthesis (*Aqp8, Cyp51, Dhcr24, Fdft1, Fdps, Hmgcs1, Idi1, Msmo1, and Sqle*). This, along with the consistent activation of cholesterol synthesis genes by the three genotoxic agents used in our study, indicates that exposure could lead to activation of SREBP2, the primary transcription factor controlling cholesterol synthesis. Indeed, research has shown that each of these three compounds can impact cholesterol homeostasis (Depass and Morris 1982; Rotimi et al. 2017; Zhang, Lu, et al. 2022). Together, this indicates that further refinement of the p53 consensus set using more diverse chemistry may improve differentiation of the p53 and cholesterol homeostasis signaling pathways.
- There was generally consistent identification of ERα activators by the ERα male and female consensus gene sets with relatively high correlations compared to the other gene sets. The known ERα activators included 17ß-estradiol, diethylstilbestrol, estriol, ethinyl estradiol, and mestranol. There were five compounds that were identified as activators of ERα using the ERα female consensus list for which there was no known evidence of ERα activation, including predictions in the Podtelezhnikov et al. study (Podtelezhnikov et al. 2020). These included 1,2-dichlorobenzene, acetamide, anastrozole, artemisinin, and methapyrilene.

**Figure 9:**
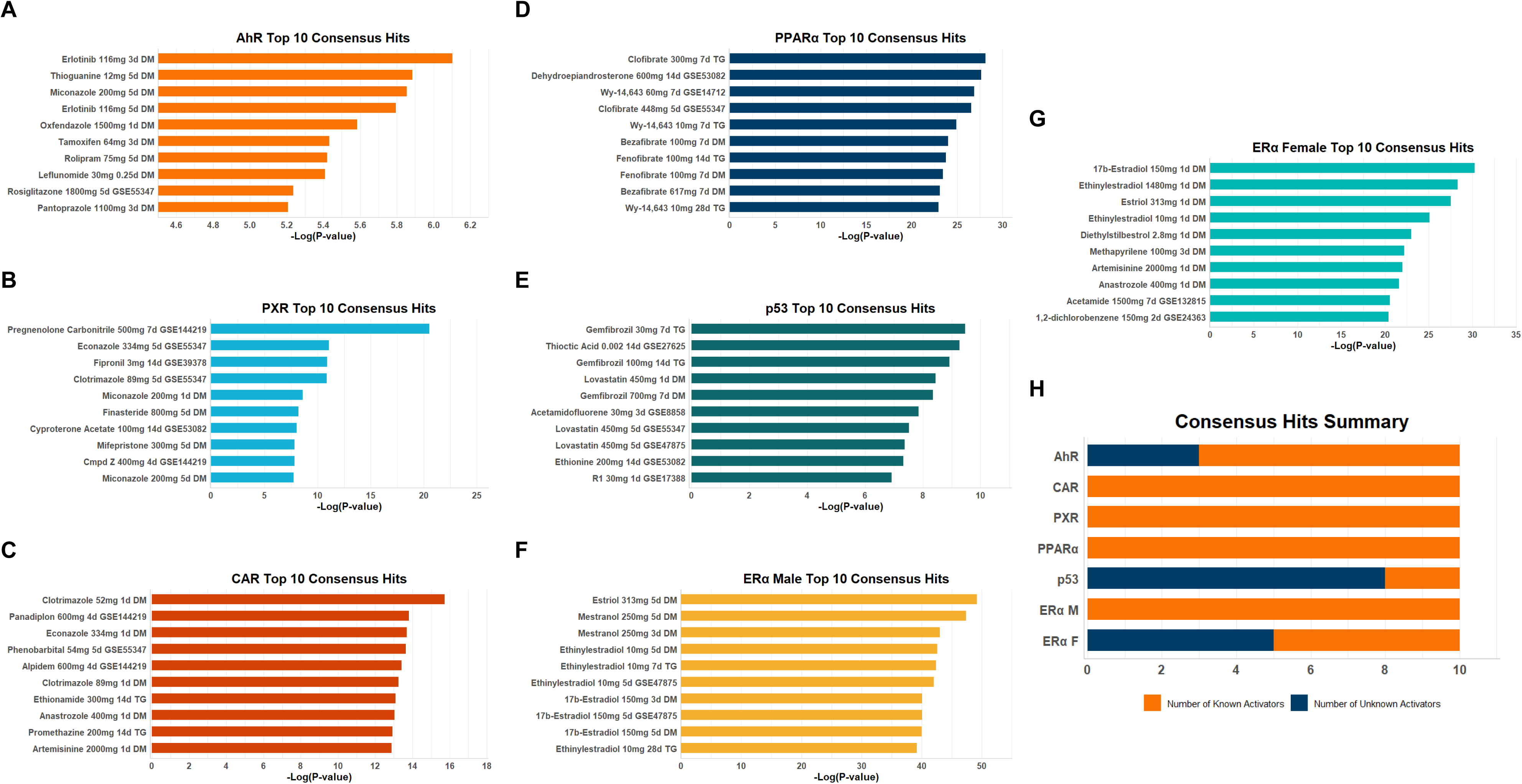
Assessing the predictive nature of TF consensus gene sets. Gene sets were identified as described in the Methods and then used to query a compendium of profiles derived from the livers of chemically treated rats. Each of the consensus gene lists was compared in a pair-wise fashion to each gene list in the rat liver compendium using the Running Fisher test. The 10 profiles with the greatest positive correlation based on the - Log(P-value) are shown for the indicated consensus gene list. (A) AhR. (B) PXR. (C) CAR. (D) PPARα. (E) p53. (F) ERα males. (G) ERα females. (H) Summary of results showing the number of compounds known or not known to be activators of the TF out of the top 10 chemicals.

In this preliminary study, the consensus gene sets show utility for detecting known or predicted activators of the specified TF in a broader compendium of rat liver studies. However, some refinement with additional studies may improve performance, particularly for the AhR and p53 sets.

## Discussion

The transcriptomic data generated by this study will serve as a foundational resource for the development of biomarkers for the MIEs of chemical carcinogenesis. These MIEs include activation of several transcription factors (TFs): ligand-activated xenobiotic receptors AhR, PXR, CAR, PPARα, and ERα, which are implicated in rodent non-genotoxic tumorigenesis, as well as the tumor suppressor protein p53, which is activated by diverse cellular stressors of potential relevance to carcinogenesis, including genotoxicity. To ensure MIE-specific transcriptional responses, we used both WT and KO rats in our studies, allowing us to identify TF-dependent genes regulation after activation of each TF. We also used three different activators for each TF at doses known to promote liver tumors in the two-year rat bioassay. Notably, this is the first comprehensive effort that has identified MIE-specific transcriptomes in rats using multiple known activators along with both WT and KO animals. While much of the early work in transcriptomic biomarker development has been conducted in mice (Furihata and Suzuki 2023; Jonker et al. 2009; Kossler et al. 2015; Park et al. 2011), rats are used heavily for agrochemical and pharmaceutical safety testing. Existing literature on rat studies has used a variety of methods, including different treatment regimes, and experimental models, encompassing variations in dose, duration, route, age, and chemical type (Corton et al. 2024; Furihata and Suzuki 2023; Furihata et al. 2020; Gong et al. 2014; Ledbetter et al. 2024; Podtelezhnikov et al. 2020; Rao et al. 2018). We employed short-term (5 days) *in vivo* assays, as they can capture early biological responses to compound exposures in a cost-effective and mechanistically informative manner, aligning with the transition away from two-year bioassays (Mitchell et al. 2025).

The TFs studied here included five nuclear receptors that function as major xenosensors in the liver (Kotulkar et al. 2024). AhR modulates the metabolism of endogenous compounds and xenobiotics, and its sustained hyperactivation is associated with the carcinogenicity of multiple environmental toxicants (Fan et al. 2024; Vogel et al. 2020). PPARα is a master regulator of hepatic lipid homeostasis (Berthier et al. 2021; Kersten and Stienstra 2017), and its activation has been identified as a major nongenotoxic mechanism of liver cancer in rodents (Klaunig et al. 2003). ERα is the primary receptor that mediates estrogenic effects in both male and female livers and is a key regulator of energy homeostasis and metabolic health (Khristi et al. 2019; Mahboobifard et al. 2022). Its expression has also been associated with sex-based differences and poor prognostic outcomes in hepatocellular carcinoma (Wang et al. 2025). CAR and PXR were historically perceived as functionally redundant due to their overlapping regulation of xenobiotic and endobiotic metabolism (Timsit and Negishi 2007). More recent studies, however, have revealed that CAR and PXR each have distinct regulatory roles in addition to coordinating functions in shared pathways. Sustained CAR activation is a well-established mechanism for rodent liver tumorigenesis, whereas the role of PXR in chemical carcinogenesis remains less clearly defined (Bwayi et al. 2022; Oladimeji et al. 2016; Yoshinari and Shizu 2022), highlighting the need to clarify the unique contributions of both receptors.

While activation of these xenobiotic receptors represents nongenotoxic mechanisms of liver toxicity, this study also assessed the transcriptional effects of p53 in response to compounds that cause DNA damage. The p53 protein is a well-documented tumor suppressor, and evidence consistently implicates its loss of function in liver tumorigenesis as well as tumorigenesis in other tissues (Link and Iwakuma 2017; Makino et al. 2022; Rahadiani et al. 2023). As p53 is activated in response to DNA damage, the expression of p53 target genes can be used as a sentinel for potential genotoxicity, although other stressors can also activate this pathway. Indeed, assays of p53 activation are frequently used as a surrogate in the assessment of genotoxicity *in vitro* (Corton, Witt, and Yauk 2019; Hsieh et al. 2019; van der Linden et al. 2014). However, when p53 activation is observed, definitive assessment of the specific MIE requires follow-up investigation, including standard genotoxicity assays. Together, the combined framework of p53 and nuclear receptor activation could be used to create a comprehensive and complementary suite of biomarkers for the detection of both non-genotoxic and potentially genotoxic MIEs of rodent liver-mediated carcinogenesis in short-term studies. Furthermore, this suite of biomarkers could inform on the potential mechanisms of neoplastic or pre-neoplastic findings in longer duration studies.

Activation of each TF by three different compounds (**Table 1**) allowed us to identify more sensitive and specific transcriptomic signatures. These gene lists were then used to curate a list of common differentially expressed genes (DEGs) specific to each TF (**Files S1a-S1g**). It is important to note, however, that some compounds produced weaker responses than others within their respective groups. This is not unexpected, as the doses used in the study were designed primarily based on tumor findings from two-year studies, whereas our experiments span only five days. For some compounds, it is possible that the tumorigenic dose in the two-year studies was based on alternative pathways to the TFs being evaluated here. Alternatively, some compounds may require additional time to produce a more pronounced effect. For instance, Cyp4a1 is a well-documented target of PPARα in rat liver and was induced by clofibrate and PFOA in this study as expected (log_2_FC = 5.2 and 5.1, respectively). Comparatively, fenofibrate produced a weaker transcriptional response (log_2_FC = 2.5). Acox1, another well-documented target of PPARα, was also induced by clofibrate and PFOA as expected (log_2_FC = 3.7 and 3.2, respectively). However, with fenofibrate, Acox1 (log_2_FC = 0.8) failed to meet the fold-change threshold. To accommodate such compound-specific differences, genes were considered if responsive to two or more of three treatments, a practice consistent with previous research (Oshida, Vasani, Jones, et al. 2015; Oshida, Vasani, Thomas, Applegate, Gonzalez, et al. 2015; Oshida, Vasani, Thomas, Applegate, Rosen, et al. 2015; Rooney, Oshida, et al. 2018).

An additional interesting finding of our studies is the interaction between PXR and CAR. To investigate the previously demonstrated crossover in transcriptomes of PXR and CAR, we employed three carefully selected activators: spironolactone, phenobarbital, and PCN in both PXR and CAR studies. PCN is well known to preferentially activate PXR whereas phenobarbital preferentially activates CAR. We selected spironolactone based on literature in mice that demonstrates it activates both CAR and PXR. However, comparison of WT and KO data showed that, at the doses and duration used in this study, spironolactone is a highly preferential agonist for rat PXR but not for rat CAR, which is a novel finding.

While our analyses identified short lists of the most robust and consistent DEGs for each TF, the biological effects of MIE activation are often mediated by coordinated changes across broader gene networks. To address this, we used Gene Set Enrichment Analysis (GSEA) to determine if expected TF-associated pathways were enriched based on the global transcriptional response to treatment. This allowed us to evaluate whether short-term exposures captured anticipated pathway-level perturbations consistent with each MIE and associated mechanisms of liver carcinogenesis, even when individual genes showed modest responses (Subramanian et al. 2005). In particular, we assessed pathways that were commonly enriched among activators within each study group. For Instance, the xenobiotic metabolism pathway was positively enriched after AhR activation, which was expected (Larigot et al. 2018). While the role of PXR in chemical carcinogenesis is less clear, enrichment of xenobiotic and fatty acid metabolism pathways was expected (Cai, Young, and Xie 2021). CAR activation expectedly enriched xenobiotic and fatty acid metabolism (Chen et al. 2019; Timsit and Negishi 2007). PPARα is well known as a key regulator of hepatic lipid homeostasis, so effects on fatty acid metabolism and peroxisome-related pathways were expected. For p53, activation of p53-enriched stress response pathways included the p53 pathway, apoptosis, and reactive oxygen species signaling (Kastenhuber and Lowe 2017; Vousden and Prives 2009). ERα is more difficult to analyze due to the sexually dimorphic context of hormone signaling in the liver. However, both males and females showed concurrent downregulation of fatty acid and bile acid metabolism, which aligns with ERα’s role in modulating hepatic metabolic gene expression (Palmisano, Zhu, and Stafford 2017; Yamamoto et al. 2006). While enrichment patterns included pathways consistent with established TF function, other significantly enriched pathways were also observed. The specificity of these findings, and whether they may be unrelated or secondary to activation of the targeted MIE, was beyond the scope of this study. However, distinguishing relevant pathways from those that are off target could be addressed with additional activators.

Past studies have also used transcript profiles generated in WT and transcription factor KO mouse livers to identify genes that exhibit consistent behavior across chemical exposures (Oshida, Vasani, Jones, et al. 2015; Oshida, Vasani, Thomas, Applegate, Gonzalez, et al. 2015; Oshida, Vasani, Thomas, Applegate, Rosen, et al. 2015). The gene expression biomarkers generated from these studies have predictive balanced accuracies that range between 91-98%. Given the success of these past studies, we carried out a preliminary analysis to determine if sets of genes with consistent behavior derived from our study could be used as potential biomarkers for identification of MIE activation in the livers of treated rats (**File S2b**). Most of the compound treatments came from the TG-GATES and DrugMatrix datasets, which include compounds that are well known activators of the MIEs under consideration. Our approach resulted in highly reliable predictions for ligand activated TFs PXR, CAR, PPARα, and ERα males. It also provided promising results for AhR and ERα in females. The predictions were not as reliable for p53 as they detected some treatments not known to induce p53. This along with the preliminary inclusion of some genes known to be modulated by other TFs, indicates that further refinement is needed to improve specificity and sensitivity of this gene set (**Figure 9a-h**).

This represents a general limitation of signature identification efforts that rely on a limited number of compounds due to their confounding effects on multiple transcriptional pathways in the liver. To this end, our Working Group has proposed a strategy to align and refine the liver MIE signatures reported here and elsewhere by adjudicating the accuracy of the included genes across an integrated set of rat liver transcriptional data. The present study will be particularly useful for these efforts as it adds experiments in knock out animals for each TF and demonstrates the requirement of each TF for the corresponding gene set. Thus, this study contributes to broader ongoing efforts to determine the predictive accuracy of rat liver TF gene sets and to qualify their usefulness as biomarkers for the screening of MIE activation or the determination of chemical mode of action.

Our studies have also opened the way to several future directions, including the determination of chemical off-target effects. For example, we used PFOA as an activator of PPARα. However, our studies using WT and PPARα-KO rats showed that PFOA, unlike fenofibrate and clofibrate, induces transcriptomic changes in the PPARα-KO rats. The genes activated by PFOA in PPARα-KO rats are mainly CAR targets, which is consistent with previous findings (Cheng and Klaassen 2008). A detailed analysis of transcriptomes activated by all chemicals in the study will yield insights into TF crosstalk. Similarly, different chemicals that activate the same TF show a significant difference in the specific genes they activate. A deeper analysis of each TF study to identify these differences will reveal additional information on sensitivity and specificity of different chemical ligands in activating TFs. Finally, a detailed analysis of control WT vs. control KO datasets will be useful in identifying basal regulation of gene expression for each of the TFs.

There is a critical need for innovation in carcinogenicity assessment driven by concerns about the translational accuracy of the two-year rodent bioassay and the mounting pressure to minimize the use of animals in research. Omics technologies, when integrated into short-term *in vivo* studies, can improve the sensitivity and mechanistic insight of toxicity testing, allowing early detection and de-risking of potential liabilities (Mitchell et al. 2025; Sewell et al. 2024; Singh et al. 2024). This integrated approach leverages the strengths of both paradigms while simultaneously enhancing the efficiency and predictive power of chemical and pharmaceutical risk assessment. This study is the first in many steps toward the primary goal of reducing the need for laborious, resource-intensive two-year rodent studies by combining a weight-of-evidence strategy with short-term *in vivo* experiments.

## Acknowledgements

The information in this document has been funded in part by the U.S. Environmental Protection Agency. The authors declare they have no actual or potential competing financial interests. This study has been subjected to review by the Office of Chemical Safety and Pollution Prevention and approved for publication. Approval does not signify that the contents reflect the views of the U.S. Environmental Protection Agency, nor does mention of trade names or commercial products constitute endorsement or recommendation for use.

## Supplemental Figure Legends

**Supplemental Figure 1:**
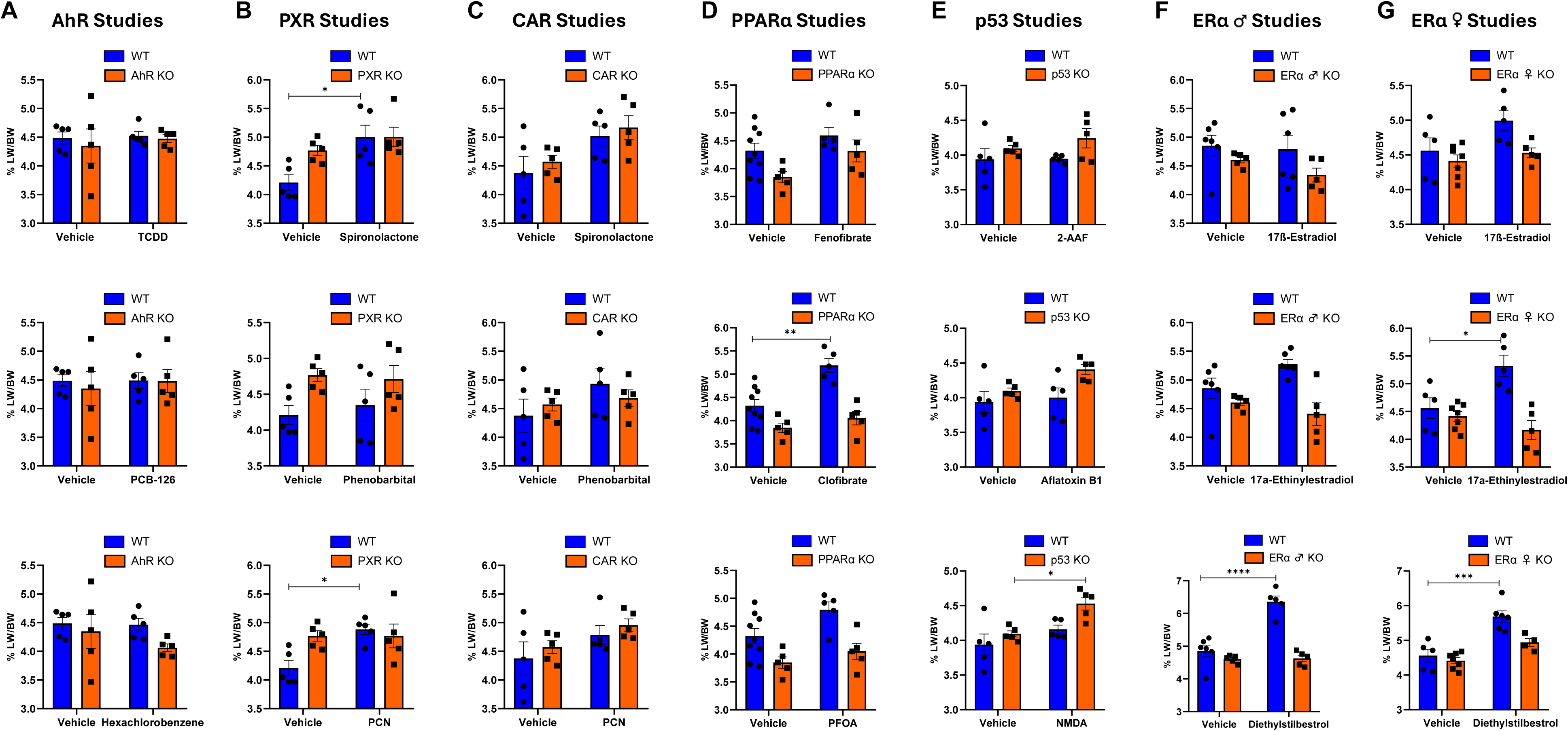
Effect of activator treatment on liver weight to body weight (LW:BW) ratio. (A) LW:BW ratio in WT and AhR KO rats after treatment with TCDD, PCB-126, or HCB. (B) LW:BW ratio in WT and CAR KO rats after treatment with Spironolactone, Phenobarbital, or PCN. (C) LW:BW ratio in WT and PXR KO rats after treatment with Spironolactone, Phenobarbital, or PCN. (D) LW:BW ratio in WT and PPARα KO rats after treatment with Fenofibrate, Clofibrate, or PFOA. (E) LW:BW ratio in WT and p53 KO rats after treatment with 2-AAF, AFB1, or NDMA. (F) LW:BW ratio in male WT and ERα KO rats after treatment with 17ß, 17α, or DES. (G) LW:BW ratio in female WT and ERα KO rats after treatment with 17ß, 17α, or DES.

**Supplemental Figure 2:**
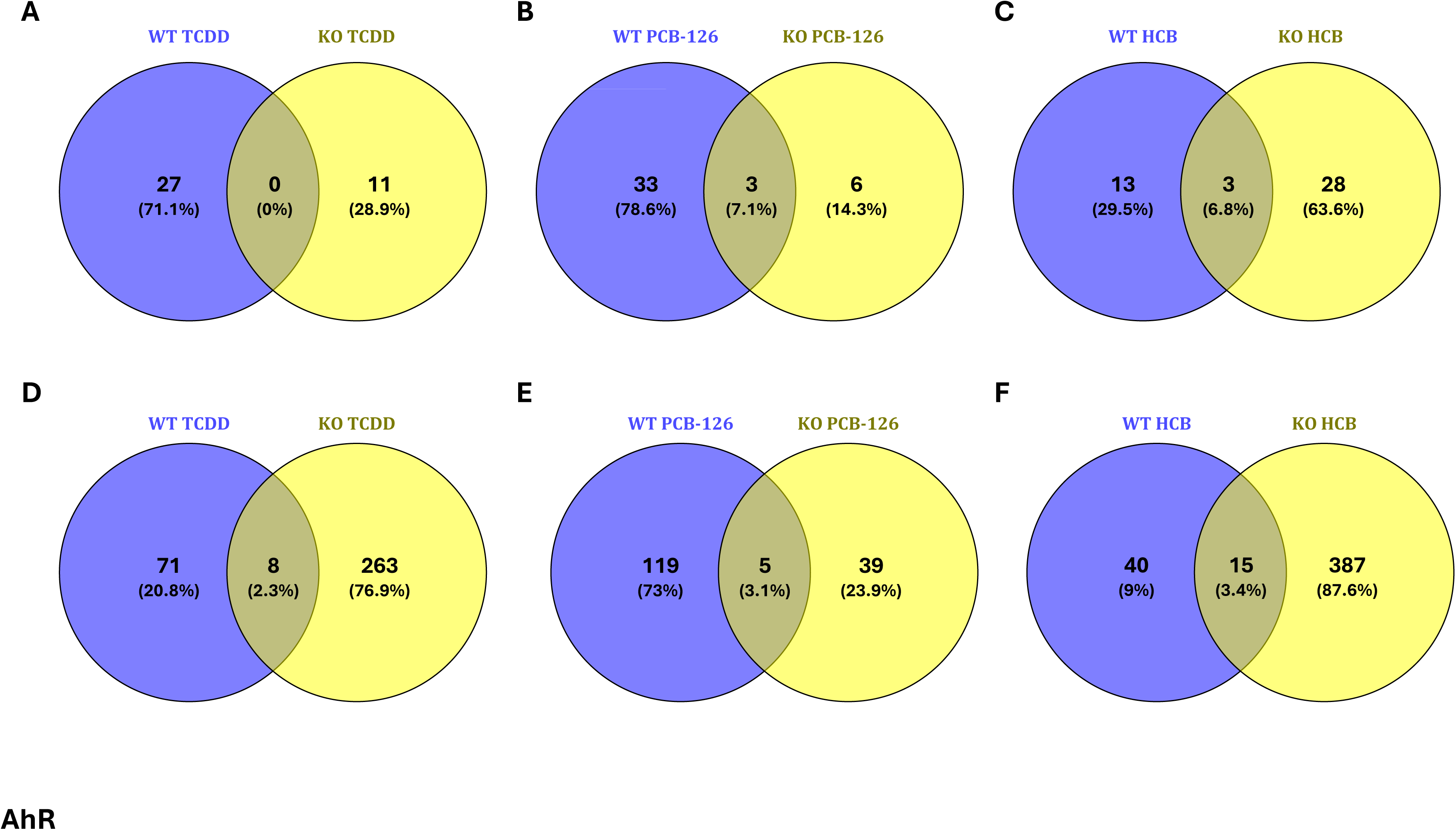
Isolation of DEGs unique to WT and AhR KO treated rats. (A) Venn diagram showing number of unique and common DEGs in WT and KO rats after TCDD treatment. (B) Venn diagram showing number of unique and common DEGs in WT and KO rats after PCB-126 treatment. (C) Venn diagram showing number of unique and common DEGs in WT and KO rats after HCB treatment. Significance threshold: p-value < 0.05 and |log_2_FC| ≥ 1. (D) Venn diagram showing number of unique and common DEGs in WT and KO rats after TDCC treatment using only p-value < 0.05. (E) Venn diagram showing number of unique and common DEGs in WT and KO rats after PCB-126 treatment using only p-value < 0.05. (F) Venn diagram showing number of unique and common DEGs in WT and KO rats after HCB treatment using only p-value < 0.05.

**Supplemental Figure 3:**
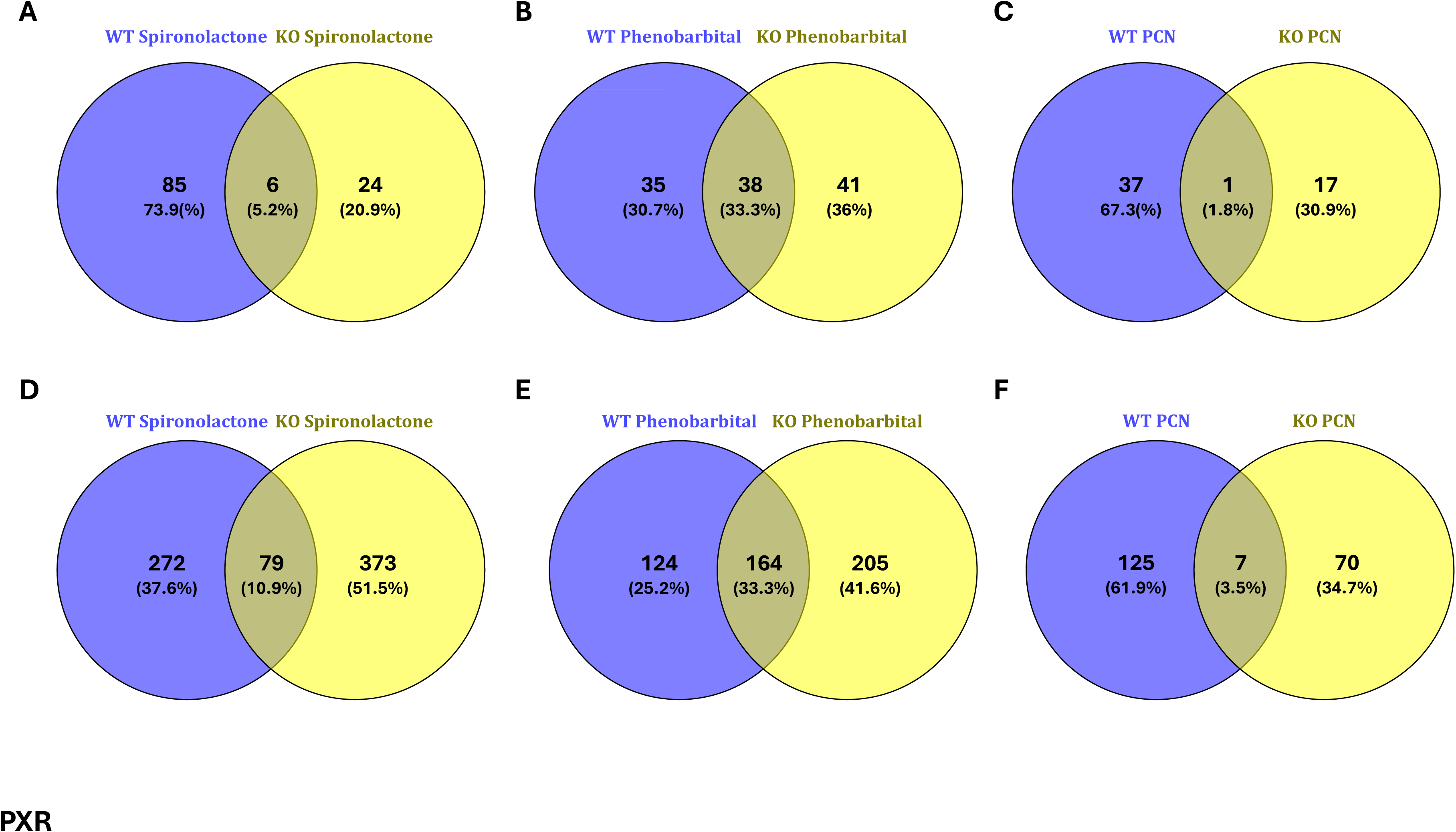
Isolation of DEGs unique to WT and PXR KO treated rats. (A) Venn diagram showing number of unique and common DEGs in WT and KO rats after Spironolactone treatment. (B) Venn diagram showing number of unique and common DEGs in WT and KO rats after Phenobarbital treatment. (C) Venn diagram showing number of unique and common DEGs in WT and KO rats after PCN treatment. Significance threshold: p-value < 0.05 and |log_2_FC| ≥ 1. (D) Venn diagram showing number of unique and common DEGs in WT and KO rats after Spironolactone treatment using only p-value < 0.05. (E) Venn diagram showing number of unique and common DEGs in WT and KO rats after Phenobarbital treatment using only p-value < 0.05. (F) Venn diagram showing number of unique and common DEGs in WT and KO rats after PCN treatment using only p-value < 0.05.

**Supplemental Figure 4:**
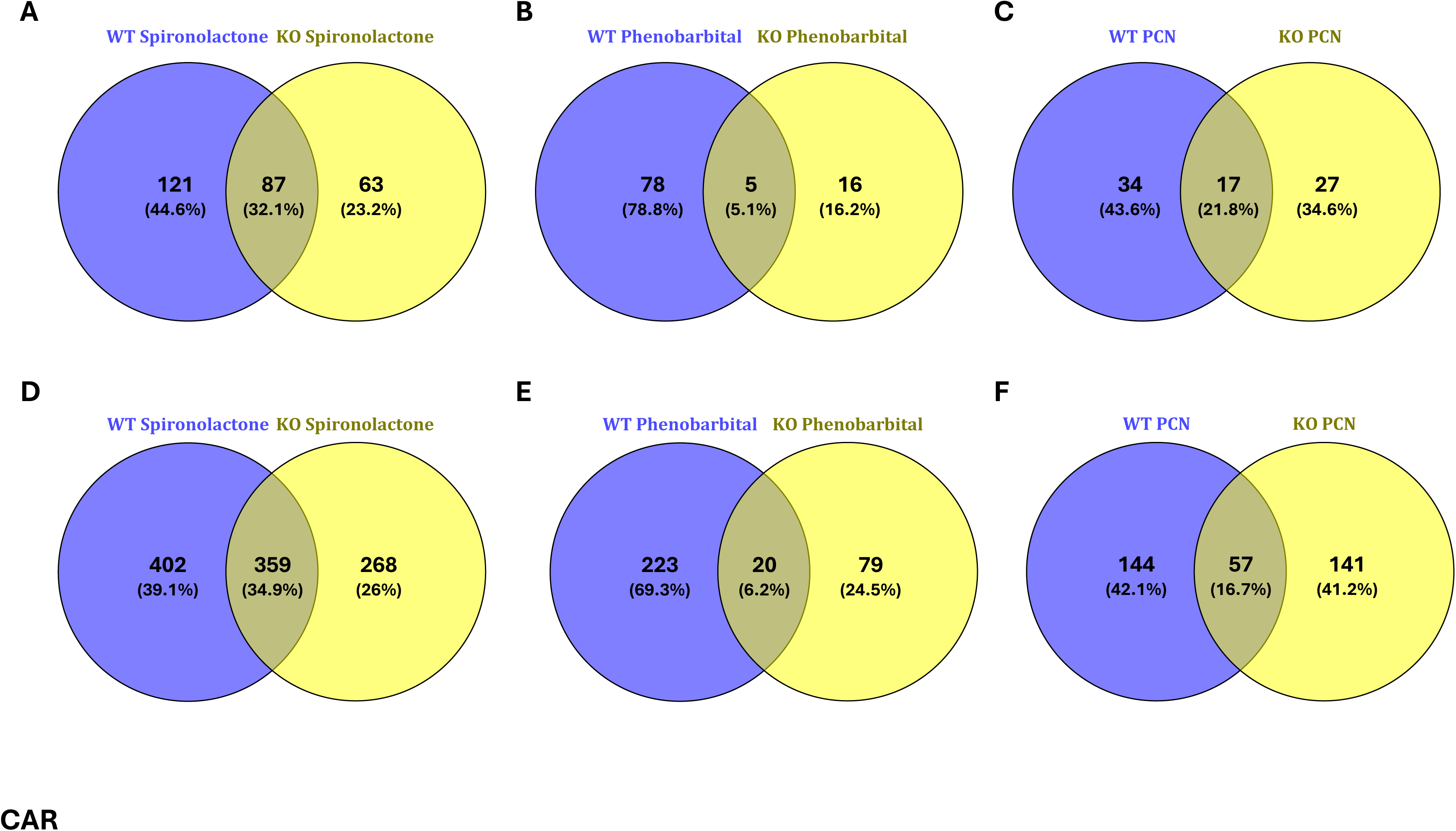
Isolation of DEGs unique to WT and CAR KO treated rats. (A) Venn diagram showing number of unique and common DEGs in WT and KO rats after Spironolactone treatment. (B) Venn diagram showing number of unique and common DEGs in WT and KO rats after Phenobarbital treatment. (C) Venn diagram showing number of unique and common DEGs in WT and KO rats after PCN treatment. Significance threshold: p-value < 0.05 and |log_2_FC| ≥ 1. (D) Venn diagram showing number of unique and common DEGs in WT and KO rats after Spironolactone treatment using only p-value < 0.05. (E) Venn diagram showing number of unique and common DEGs in WT and KO rats after Phenobarbital treatment using only p-value < 0.05. (F) Venn diagram showing number of unique and common DEGs in WT and KO rats after PCN treatment using only p-value < 0.05.

**Supplemental Figure 5:**
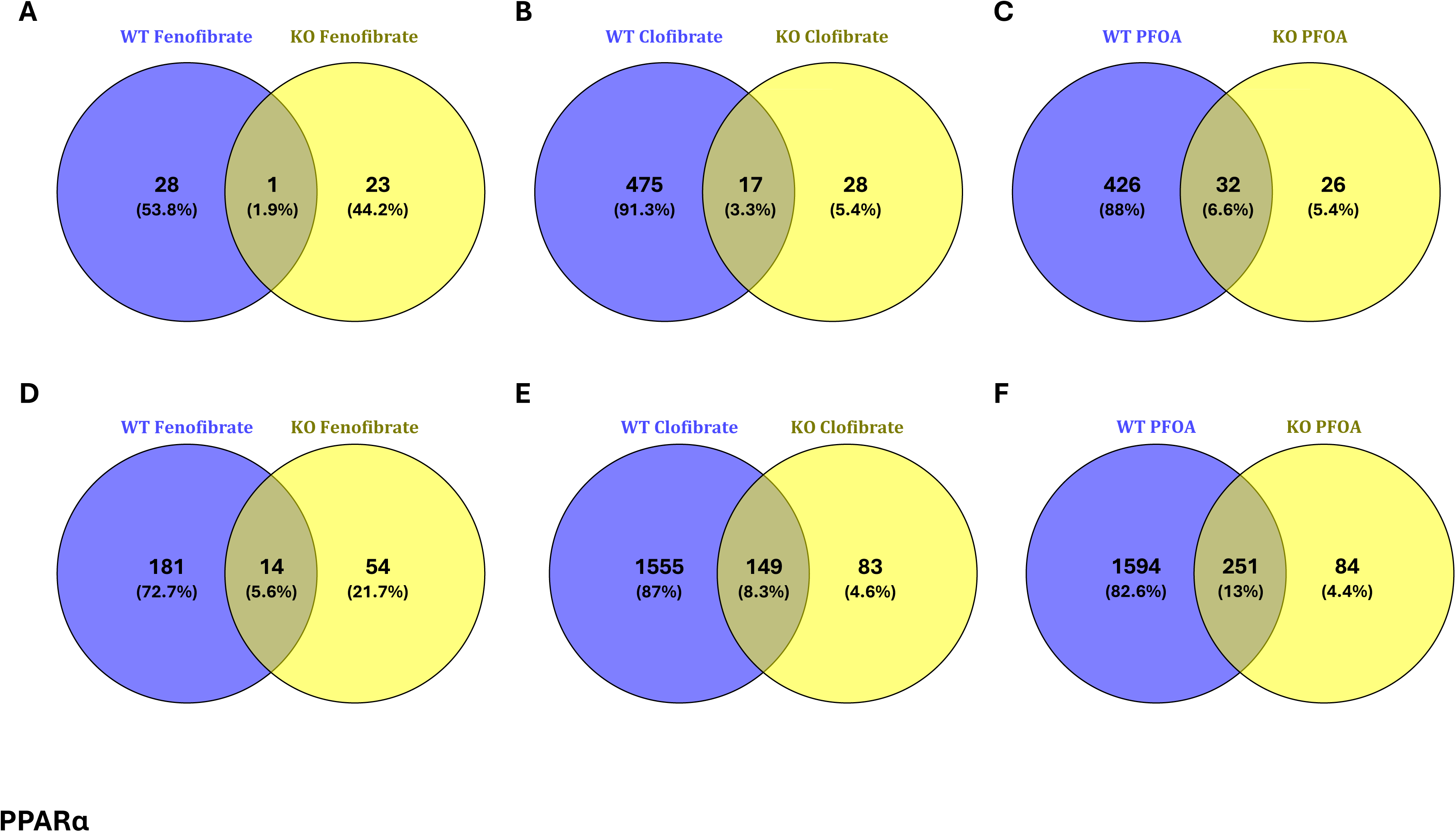
Isolation of DEGs unique to WT and PPARα KO treated rats. (A) Venn diagram showing number of unique and common DEGs in WT and KO rats after Fenofibrate treatment. (B) Venn diagram showing number of unique and common DEGs in WT and KO rats after Clofibrate treatment. (C) Venn diagram showing number of unique and common DEGs in WT and KO rats after PFOA treatment. Significance threshold: p-value < 0.05 and |log_2_FC| ≥ 1. (D) Venn diagram showing number of unique and common DEGs in WT and KO rats after Fenofibrate treatment using only p-value < 0.05. (E) Venn diagram showing number of unique and common DEGs in WT and KO rats after Clofibrate treatment using only p-value < 0.05. (F) Venn diagram showing number of unique and common DEGs in WT and KO rats after PFOA treatment using only p-value < 0.05.

**Supplemental Figure 6:**
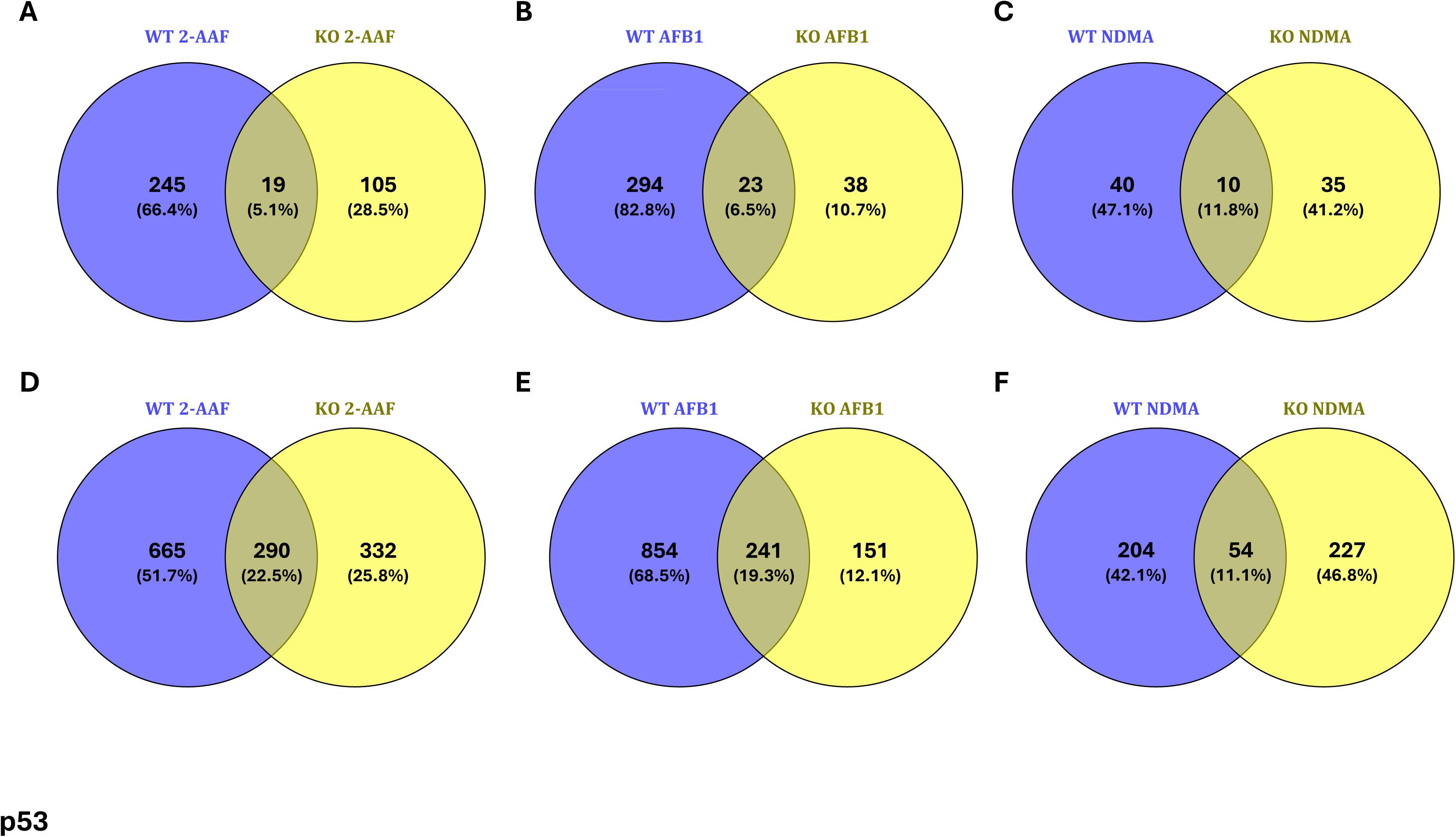
Isolation of DEGs unique to WT and p53 KO treated rats. (A) Venn diagram showing number of unique and common DEGs in WT and KO rats after 2-AAF treatment. (B) Venn diagram showing number of unique and common DEGs in WT and KO rats after AFB1 treatment. (C) Venn diagram showing number of unique and common DEGs in WT and KO rats after NDMA treatment. Significance threshold: p-value < 0.05 and |log_2_FC| ≥ 1. (D) Venn diagram showing number of unique and common DEGs in WT and KO rats after 2-AAF treatment using only p-value < 0.05. (E) Venn diagram showing number of unique and common DEGs in WT and KO rats after AFB1 treatment using only p-value < 0.05. (F) Venn diagram showing number of unique and common DEGs in WT and KO rats after NDMA treatment using only p-value < 0.05.

**Supplemental Figure 7:**
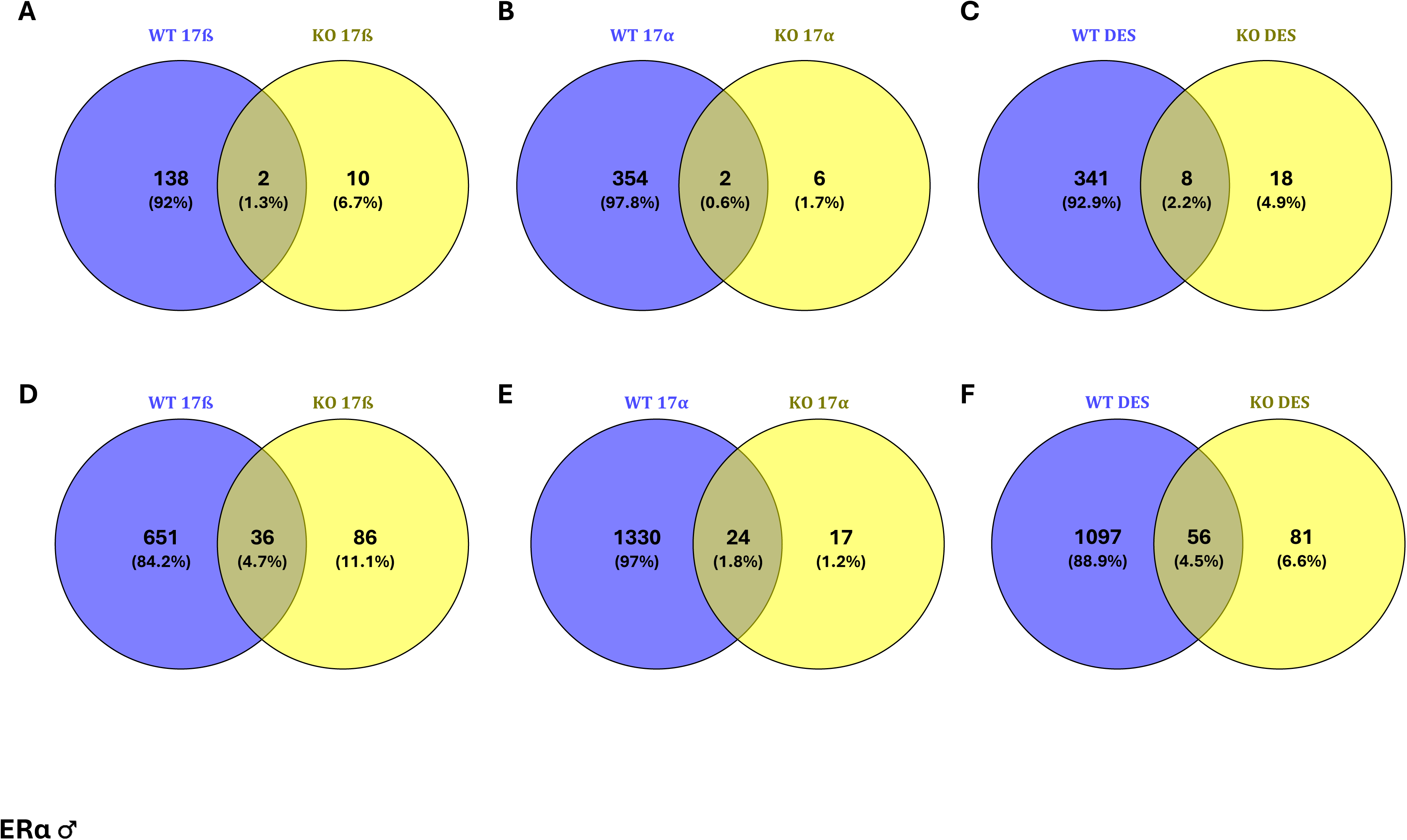
Isolation of DEGs unique to WT and ERα male KO treated rats. (A) Venn diagram showing number of unique and common DEGs in WT and KO rats after 17ß treatment. (B) Venn diagram showing number of unique and common DEGs in WT and KO rats after 17α treatment. (C) Venn diagram showing number of unique and common DEGs in WT and KO rats after DES treatment. Significance threshold: p-value < 0.05 and |log_2_FC| ≥ 1. (D) Venn diagram showing number of unique and common DEGs in WT and KO rats after 17ß treatment using only p-value < 0.05. (E) Venn diagram showing number of unique and common DEGs in WT and KO rats after 17α treatment using only p-value < 0.05. (F) Venn diagram showing number of unique and common DEGs in WT and KO rats after DES treatment using only p-value < 0.05.

**Supplemental Figure 8:**
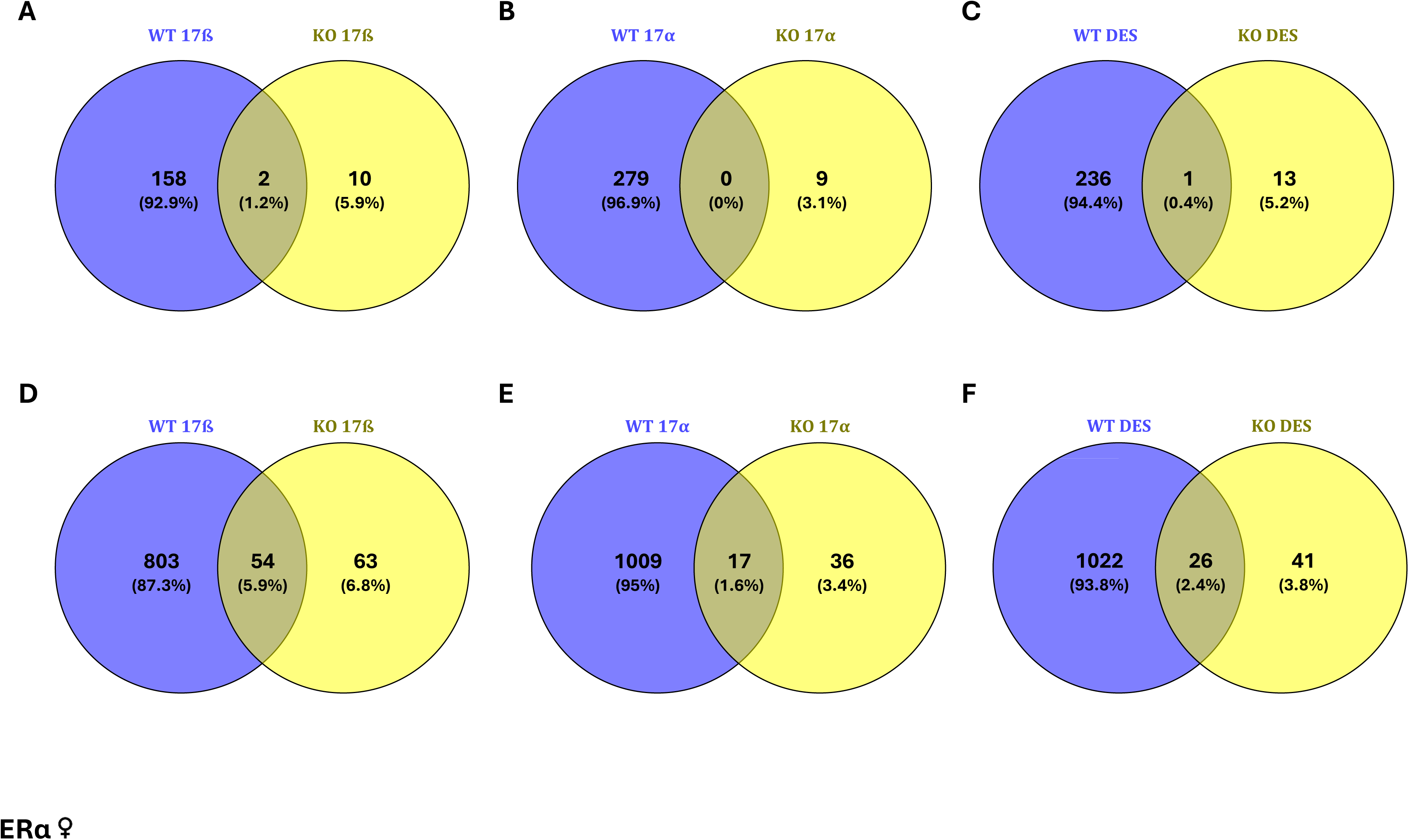
Isolation of DEGs unique to WT and ERα female KO treated rats. (A) Venn diagram showing number of unique and common DEGs in WT and KO rats after 17ß treatment. (B) Venn diagram showing number of unique and common DEGs in WT and KO rats after 17α treatment. (C) Venn diagram showing number of unique and common DEGs in WT and KO rats after DES treatment. Significance threshold: p-value < 0.05 and |log_2_FC| ≥ 1. (D) Venn diagram showing number of unique and common DEGs in WT and KO rats after 17ß treatment using only p-value < 0.05. (E) Venn diagram showing number of unique and common DEGs in WT and KO rats after 17α treatment using only p-value < 0.05. (F) Venn diagram showing number of unique and common DEGs in WT and KO rats after DES treatment using only p-value < 0.05.

## Supplemental Files

**Supplementary File 1a:** AhR DESeq2 outputs and shared DEGs.

**Supplementary File 1b:** PXR DESeq2 outputs and shared DEGs.

**Supplementary File 1c:** CAR DESeq2 outputs and shared DEGs.

**Supplementary File 1d:** PPARα DESeq2 outputs and shared DEGs.

**Supplementary File 1e:** p53 DESeq2 outputs and shared DEGs.

**Supplementary File 1f:** ERα male DESeq2 outputs and shared DEGs.

**Supplementary File 1g:** ERα female DESeq2 outputs and shared DEGs.

**Supplementary File 2a:** Consensus gene sets for all studies.

**Supplementary File 2b:** Consensus gene sets top 10 hits.

